# Seed plant families with diverse mycorrhizal states have higher diversification rates

**DOI:** 10.1101/824441

**Authors:** María Isabel Mujica, Gustavo Burin, María Fernanda Pérez, Tiago Quental

## Abstract

A crucial innovation in plant evolution was the association with soil fungi during land colonization. Today, this symbiotic interaction is present in most plants species and can be classified in four types: arbuscular (AM), Ecto (EM), Orchid (OM) and Ericoid Mycorrhiza (ER). Since the AM ancestral state, some plants lineages have switched partner (EM, OM and ER) or lost the association (no-association: NM). Evolutionary transitions to a novel mycorrhizal state (MS) might allow plant lineages to access new resources, enhancing diversification rates. However, some clades are not restricted to one MS, and this variability might promote diversification. In this study we address the relationship between MS and diversification rates of seed plant families. For this, we used the recently published FungalRoot database, which compiled data for 14,870 species and their mycorrhizal partners. We assigned a MS to each plant family, calculated the MS heterogeneity and estimated their diversification rates using the method-of-moments. Families with mixed MS had the highest diversification rates and there was a positive relationship between MS heterogeneity and diversification rates. These results support the hypothesis that MS lability promotes diversification and highlight the importance of the association with soil fungi for the diversification of plants.

## Introduction

Understanding the basis of the exceptional plant diversity has been a matter of interest for ecologist and evolutionary biologist since Darwin. Great focus has been placed on estimating plants diversification rates and identifying the factors that could influence them (Eriksson and Bremer 1992; Moore and Donoghue 2007; O’Meara et al. 2016; Vamosi et al. 2018). The acquisition of novel traits (sometimes referred to as “key innovations”), such as pollination by animals (Eriksson and Bremer 1992) or physiological seed dormancy (Willis et al. 2014), have been proposed to promote diversification of plant lineages. This “key innovation” perspective suggests that the acquisition of a novel trait might allow a given lineage to exploit the environment in a significantly different way, potentially resulting in an explosive radiation. A crucial innovation in plant evolution was the association with soil fungi during land colonization (Pirozynski and Malloch 1975; Selosse and Le Tacon 1998; Strullu-Derrien et al. 2018). Before plant colonization, land was hostile, with extreme drought and temperatures, and barren rocky substrate; hence, the association with terrestrial fungi allowed the algae ancestors of plants to successfully colonize the land (Selosse et al. 2015). This initial symbiotic association was the prelude of modern mycorrhizas (Feijen et al. 2018), the association between fungi and root plants in which plants transfer carbon to fungi and receive nutrients in return (Smith and Read 2008). Today, this symbiosis is present in 86% of land plants species (Heijden et al. 2015) and based on their structure and function can be classified in four major types: arbuscular mycorrhiza (AM), ectomycorrhiza (EM), orchid mycorrhizal (OM) and ericoid mycorrhizal (ER) (Brundrett 2002).

Ancestral state reconstruction and the fossil record suggest that the ancestor of seed plants probably had AM associations (Redecker et al. 2000; Maherali et al. 2016). This is the most frequent mycorrhizal type in plants (74% of extant plant species) and is characterized by an association with Glomeromycete fungi (Heijden et al. 2015). Between 100 and 200 million years ago, some lineages switched fungal partners to several lineages of Basidiomycetes, forming what is described as the EM associations (Brundrett 2002). The acquisition of EM resulted in new root functional capabilities as freezing tolerance (Lehto et al. 2008), which seem related to the dominance of EM angiosperms and gymnosperm in cool forests (Brundrett 2002). Similarly, Orchidaceae and species within the Ericaceae family recruited new fungal lineages and formed OM and ER associations respectively. Orchids associate with fungal families Ceratobasidiaceae, Tulasnellaceae and Sebacinaceae, which in addition to nutrient exchange, promote seed germination which cannot germinate without mycorrhizal support (Rasmussen 2002). Ericoid mycorrhizal associations (ER), on the other hand, involve mainly fungi from Sebacinales and Helotiales and are mostly frequent under acidic and infertile heathland conditions (Perotto et al. 2002; Heijden et al. 2015). Finally, some lineages have lost their mycorrhizal associations and became non-mycorrhizal (NM). This transition has frequently occurred through an intermediate state of facultative arbuscular mycorrhiza (AM) plants (Maherali et al. 2016). Some of NM lineages evolved alternative resource-acquisition strategies (Werner et al. 2018) like cluster-roots in Proteaceae (Neumann and Martinoia 2002) or parasitism in Loranthaceae (Wilson and Calvin 2006). Therefore, since the AM ancestral state some plant lineages have followed different mycorrhizal evolutionary pathways: completely switched partner (EM, OM and ER) or lost the association (Werner et al. 2018). Evolutionary transitions to a novel mycorrhizal state might allow plant lineages to access unexplored ecological resources, facilitating them to colonize environments that were not available before, and possibly enhancing their diversification rates. However, there are lineages in which some species acquire a new mycorrhizal state and at the same time, other species retain the ancestral state (AM) (Brundrett, 2002) increasing the variability of mycorrhizal states, which might in fact promote diversification of these lineages.

Although there are several traits known to influence plant diversification (Vamosi et al 2018; Hernandez & Wiens 2020), and mycorrhizal symbiosis has been pointed out as a key factor in the evolution and diversification of land plants (Brundrett and Tedersoo 2018; Feijen et al. 2018) the effect of mycorrhizal symbiosis on plant diversification has not been evaluated before. Hence the hypotheses that switching or adding new partners might increase diversification rates of plant lineages have not been formally evaluated. The few studies available from the fungal perspective suggest that shifts in mycorrhizal associations might affect diversification of involved partners (Sánchez-García and Matheny 2017; Sato et al. 2017).

Our aim was to specifically evaluate whether and how mycorrhizal associations may affect plant diversification dynamics rather than to identify all possible traits that might have promoted the diversification of plants. For this, we address the following questions: (1) Do the lineages that established derived mycorrhizal associations present higher diversification rates than the ones that retained the ancestral mycorrhizal state? This investigates the idea of a key innovation mechanism of diversification; and (2) Is there a relationship between mycorrhizal variability and diversification rates among different plant lineages? This would investigate the idea that evolutionary lability might increase diversification dynamics. To answer these questions, we explored the relationship between the mycorrhizal state and the diversification rates of most of seed plant families.

## Materials and Methods

### Mycorrhizal state database

To obtain information of plant species and their mycorrhizal state, we used the FungalRoot database, a recently published global databank of plant mycorrhizal associations (Soudzilovskaia et al. 2019). This database compiles previous lists and surveys of plant species and their mycorrhizal associations, including 36,303 records for 14,870 plant species. Based on these empirical records and on expert opinion, the authors proposed a list of mycorrhizal status at the plant genus level, which contains 14,541 total genera, from which 12,558 correspond to seed plant genera, that results in information for 295,221 seed plant species. Given that it comprises information for a much larger number of species, we used the genus-level list for our main analyses, but we note it assumes no variation within genera. An alternative dataset constructed directly at the species level (representing a smaller dataset) was also used (see below) and their results, which are congruent to our main analysis, are presented at the supplementary information.

The genus-level list from FungalRoot database includes information for genera belonging to 413 seed plant families. In this list, genera were classified as AM, EM, NM, OM, ER, or with multiple mycorrhizal status (i.e. AM-EM, AM-NM, named MIX genera). From those families, 33 families were discarded because either they had their entire richness classified as unknown type (21), had no data on stem age (7), represented states with no or few evolutionary replicas (3 - namely Orchidaceae, Ericaceae and Diapensiaceae), or for being represented by two different tips in the phylogeny (2). We then used a base dataset containing 380 plant families for the entirety of the paper.

### Addressing caveats and uncertainties in the Database

We used the FungalRoot database to assign a unique mycorrhizal state to each seed plant family and to calculate the proportion of mycorrhizal states within them (see below). However, this database presents some uncertainties about plants mycorrhizal states that were addressed to mitigate any potential biases that could affect the classification of families.

First, for genera with multiple mycorrhizal status (MIX genera) there is no information about how mycorrhizal states are distributed among its constituent species. Instead of using multiple mycorrhizal status as a separate state, we divided the species richness of those genera into the mycorrhizal states that composed its category (e.g. AM-NM was divided into AM and NM states). To avoid possible biases derived from choosing an arbitrary proportion of species for each state and also to properly address the uncertainty regarding the proportions of species in each category for MIX genera, we generated 10000 datasets, each of which contained a random division of the total richness of each MIX genus among their corresponding unique types (e.g. if one genus had 100 species and was classified originally as AM-NM, we would sample a number between 1 and 100 and then assign this sampled number of species to the AM category and the remaining species to the NM category prior to grouping all data per family). We obtained the richness of each genus from The Plant List ([theplantlist.org]), and then calculated the number of species with each mycorrhizal state within each family, discarding those genera that had unknown mycorrhizal type. All steps described below were applied individually to each of the 10000 datasets.

Second, Brundrett & Tedersoo (2019) pointed out potential mistakes in mycorrhizal type identification on large databases, and how these misdiagnoses might lead to wrong conclusions. Although the approach used by these authors to determine these errors (taxonomic approach; Brundrett (2017)) is controversial (Bueno et al. 2019; Sun et al. 2019), and the proportion of errors they detected in databases is relatively low (Brundrett and Tedersoo 2019), Brundrett and Tedersoo (2019) are right to point out that caution must be taken when analyzing large databases of plant mycorrhizal status. Therefore, to assess the effect of mycorrhizal misassignment in the genus-level dataset, we introduced errors to the mycorrhizal state on 20% of plant species (one order of magnitude higher than the error estimated from Brundrett and Tedersoo (2019)). This was done by randomly choosing 52826 species (corresponding to 20% of total plant richness) and reassigning them a new mycorrhizal state (that was assured to be different than the original). We replicated this procedure 10 times for each of the 10000 datasets used in the main analysis resulting in 100000 random datasets that were then submitted to the same analysis protocol of the main analysis. The results obtained with the “mis assigned databases” were similar to those derived from original data (Appendix S1 in Supporting Information).

Third, we evaluated if the results derived from the genus-based dataset were maintained when using only empirical data directly collected from the species-level list, which has considerably less species, but more precise information. Additionally, in the species-level list, the authors included remarks for 3,954 plant records (out of 36,303), indicating potential mistakes or misidentification of mycorrhizal associations in the original publication (see details in Table “Media 3” in Soudzilovskaia et al. (2019)). Then, to test the effect of potential errors in the species database, we conducted the analyses (i) excluding and (ii) without excluding plant records at the species level that had remarks (see Appendix S2).

### Family mycorrhizal state and diversity

For each of the 10000 datasets, each family was assigned with a unique mycorrhizal state (AM, EM, NM, ER or OM) when more than 60% of species sampled belonged to this mycorrhizal state. If no single state were present in more than 60% of species, the family was assigned as “MIX”, to indicate no dominance of any mycorrhizal association. Other thresholds for the assignment of family mycorrhizal state were tested and the pattern was similar (50%, 80% and 100%, Appendix S3). To investigate the effect of mycorrhizal diversity in the diversification dynamics we estimated the “Mycorrhizal State Diversity Index” (MSDI), which does not depend on our categorical criteria of mycorrhizal state assignment and is calculated by estimating the heterogeneity of the mycorrhizal states in each family using the Shannon diversity index. MSDI was calculated independently for each family using each of the 10000 datasets, given the uncertainty associated with genera that classified as “MIX” (see above).

### Diversification rates

First, to explore existence of rate heterogeneity among families of plant seed, we assessed the association between age and richness among seed plant families. Thus, following Sánchez-Reyes et al. (2017), we evaluated the association between stem age and richness, including all seed plant families available (i.e. without removing families lacking information on mycorrhizal states) and correcting for phylogenetic structure and not. Stem group ages of the families were obtained from the dated molecular phylogeny of seed plants of Zanne et al. (2014) and the number of species of each family was obtained from The Plant List ([http://www.theplantlist.org]) as previously mentioned. No association was found between stem group age and richness, either considering or not phylogenetic structure (R^2^=0.009; R^2^=0.007, respectively; Fig.S1), suggesting that diversification rates significantly vary among clades (Sánchez-Reyes et al. 2017) which has been demonstrated also in several studies (Igea et al. 2017; Diaz et al. 2019; Magallón et al. 2019) and thus justifies further investigation to explain such heterogeneity. Diversification rates for each seed plant family were estimated using the method-of-moments from Magallón & Sanderson (2001) and stem group ages. We are aware of other methods to investigate the association between trait states and diversification dynamics (e.g. BAMM; Rabosky, (2014)), however most of them would be unusable here, given that they require detailed species-level phylogenies and the most complete resolved seed plant phylogeny is massively undersampled at this level (it has molecular data for less than 23% of species, Smith & Brown, 2018). Because the relative contribution of extinction is unknown, we used two distinct scenarios to characterize the relative extinction rates (□) in the estimation of diversification rates, one with no extinction, □ = 0.0 and another with high extinction, □ = 0.9.

The seed plant phylogeny used in this study (Zanne et al. 2014) was built using a Maximum Likelihood framework, and therefore consists in a single topology, not allowing phylogenetic uncertainty to be readily incorporated. That said, the uncertainty of relationships among lineages at the scale we used (relationship between families) should not influence the results as much as on where the uncertainty at the species level (which might be a lot higher) is a crucial aspect. More importantly, we replicated the same procedure using a different family-level phylogeny (Harris and Davies 2016), allowing some level of phylogenetic uncertainty to be addressed. We used their phylogeny and rate estimates to run a similar analysis. Their estimates of diversification rates used the same source of information for species richness per family (theplantlist.org), but their stem age estimates for plant families are different. This results in slightly different rate estimates from our estimates. As expected, the results are virtually identical (Appendix 4) and here we show only the results using the phylogeny from the Zanne et al. (2014).

### Phylogenetic signal

To evaluate if we need to correct for phylogeny, we calculated the phylogenetic signal of mycorrhizal traits and diversification rates from the pruned phylogeny of seed plants. For this, the seed plant phylogeny (Zanne et al. 2014) was pruned to obtain a family level phylogeny, with one species per family as tips. For the continuous variables - MSDI and diversification rates - we calculated phylogenetic signal using Pagel’s Lambda (Pagel 1999) using the function phylosig in the package phytools in R (Revell 2012). For the categorical variable, mycorrhizal state, we estimated the phylogenetic signal using the D parameter (Fritz and Purvis 2010) with the function phylo.d in caper package in R (Orme et al. 2018). In the case of MSDI and mycorrhizal states, phylogenetic signal was calculated using the 10000 dataset, thus a distribution of lambda values and D parameter were obtained for each variable (see Appendix 5).

### Statistical analysis

Using the 10000 datasets of plant mycorrhizal states (see above), we performed two main analyses: (1) ANOVAs to test for potential differences in diversification rates among plant families with different mycorrhizal states and (2) regressions to evaluate the relationship between mycorrhizal diversity (MDSI) and diversification rates. Since some (but not all) of the mycorrhizal traits and diversification rates showed significant phylogenetic signal (Appendix S5), we evaluated the effect of mycorrhizal associations on diversification rates by considering and not the phylogenetic structure in the residuals. Then, for the first analysis we used both ANOVA and phylogenetic ANOVA using the function phylanova from phytools in R, where mycorrhizal state was used as factor and diversification rate as response variable. Because the mycorrhizal states OR and ER only had one and two family respectively, those were removed from this analysis. Likewise, in the second analysis we performed a linear model and a PGLS regression in the R package caper (Orme et al. 2018), with diversification rates as response variable and mycorrhizal heterogeneity as explanatory variable. For the PGLS models we used each lambda value obtained from respective previous phylogenetic signal analysis. To further explore the potential confounding effect on the association between mycorrhizal association and diversification dynamics, we performed PGLS regressions to assess the relationship between mycorrhizal diversity index, age and species richness. Given that all these analyses were performed 10000 times (using each dataset of mycorrhizal states), we obtained a distribution of p-values and R^2^.

## Results

According to our classification, and using 60% threshold for mycorrhizal state assignment, from the 380 families used in the analyses, a median of 270 families were AM (for example, Amaryllidaceae, Asteraceae and Lamiaceae), 10 were EM (like Fagaceae, Nothofagaceae, Betulaceae and Pinaceae), 56 were NM (such as Brassicaceae, Caryophyllaceae and Juncaginaceae) and 26 were mixed (Table 1 and Fig. 1). Mixed families contain species that retained the ancestral state (AM) and species that present a different mycorrhizal state (EM or NM). The phylogenetic signal strength differs among mycorrhizal states, but all mycorrhizal states are phylogenetically clustered to some extent (Fig S21). Likewise, the phylogenetic signal of diversification rates was significantly different from a random structure in in r^□ = 0.0^ and r^□ = 0.9^ (Table S17). Therefore, statistical analyses were performed correcting and not for phylogeny.

**Table 1:** Median, minimum and maximum number of families with each mycorrhizal state in the 10000 datasets. The values were calculated with the classification using a 60% threshold.

|  | AM | EM | ER | MIX | NM | OM |
| --- | --- | --- | --- | --- | --- | --- |
| Minimum | 260 | 9 | 1 | 13 | 46 | 1 |
| Median | 270 | 10 | 1 | 26 | 56 | 1 |
| Maximum | 281 | 13 | 1 | 38 | 69 | 1 |

**Figure 1.**
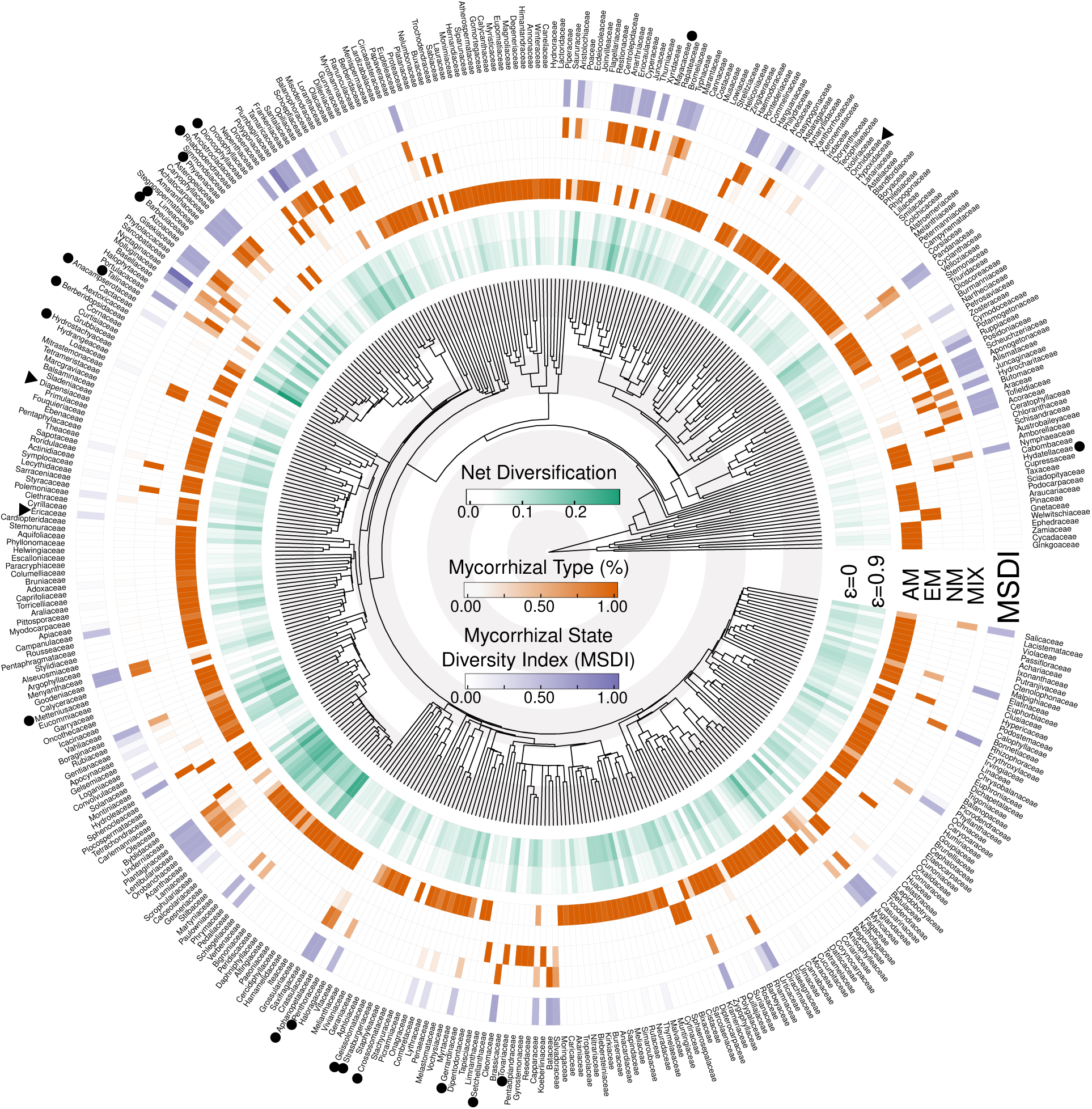
Family-level, time-calibrated phylogeny for the 405 seed plant families (based on Zanne et al 2014) and mycorrhizal data from Fungal Root database. Families with symbols were excluded from main analyses due to low or lack of mycorrhizal state replication (▴, three families: one OM and two ER) or unknown mycorrhizal states (●, N=21). For each family, the proportion of species within each mycorrhizal state is represented in the rose-to-red boxes, AM: Arbuscular mycorrhiza, EM: Ectomycorrhiza, NM: non-mycorrhizal and MIX: multiple mycorrhizal states. The mycorrhizal state diversity index (MSDI) is represented in the purple boxes and the diversification rates (r) are shown in the green boxes, for both extinction scenarios. To illustrate the timescale of the phylogeny, the width of concentric white and gray circles represents 50 million years.

When using the 60% threshold, there was a significant difference in diversification rates among mycorrhizal states (Fig.2A), which is observed in the distribution of p-values of 10000 standard ANOVAs and phylogenetic ANOVAs (median of p-values were 0.000137 and 0.018 respectively; Fig. 2B). These differences are maintained with □ (relative extinction rates) = 0.0 (median p-value=0.000109 of standard ANOVAs; median p-value=0.016 of phylogenetic ANOVAs; Fig.S10). The *a posteriori* analysis of the ANOVA showed that diversification of MIX families was significantly higher than that of AM and NM families (Fig. 2C, Table S14) and the same tendency is observed when correcting for the phylogenetic structure (Fig. 2C, Table S13). Sensitive analyses to explore the effect of the threshold to define the different categories suggest that increasing the threshold (80% or 100%) results in an even stronger difference between the MIX families and the other categories (Figs S15 and S16), while a less stringent categorization (only 50% of a certain kind would suffice to define it) results in non-significant differences when one takes into account phylogeny (Figs S9 and S13).

**Figure 2.**
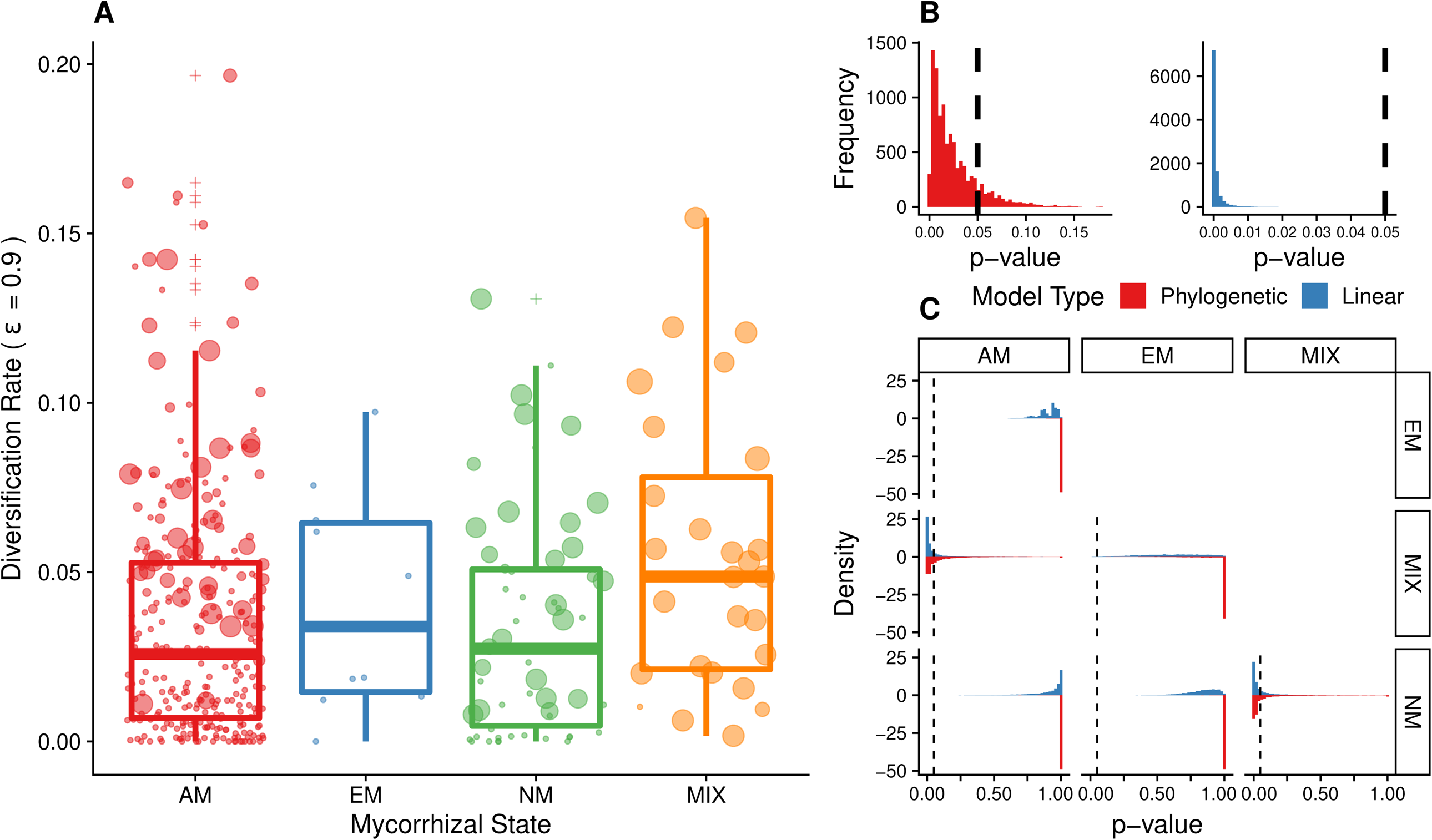
Relationship between mycorrhizal state and diversification rates using 10000 datasets of seed plant mycorrhizal states. a) Boxplot using one dataset of mycorrhizal states as example to show differences on diversification rates among families with different mycorrhizal states. Diversification rate was estimated with □ (relative extinction fraction) = 0.9. AM: Arbuscular mycorrhiza, EM: Ectomycorrhiza, NM: non-mycorrhizal and MIX (families with no dominance of any specific mycorrhizal association). The size of the points indicates the Mycorrhizal State Diversity Index (MSDI) value for each family. Panels b and c show the frequency distribution of p-values for (b) the ANOVAs and (c) the *a posteriori* comparisons among mycorrhizal states. The black dashed lines show the 0.05 value, red histograms correspond to results of phylogenetic ANOVAs and blue histograms correspond to standard ANOVAs.

The higher values of mycorrhizal state diversity index (MSDI) were found in Nyctaginaceae (median=1.09), Polygonaceae (0.98) and Rhizophoraceae (0.726), while the lowest value was zero and it was observed in 275 families that have all species in the same mycorrhizal state, like in Pinaceae (EM, n = 255), Araucariaceae (AM, n = 38) and Droseraceae (NM, n = 189). There was a positive correlation between mycorrhizal diversity index and diversification rates, observed under the two scenarios of extinction (r^□ = 0.9^ is shown in Figure 3 and with r^□ = 0.0^ is shown in Figure S17) both for the linear models and the PGLS (Fig.3B and 3C). The median of R^2^ for linear models was 0.1188 and for PGLS was 0.105, whereas the median of p-values was around 1×10^−12^ and 1×10^−11^ respectively. MSDI had no correlation with age and a significant but very low correlation with species richness (Fig. S2).

**Figure 3.**
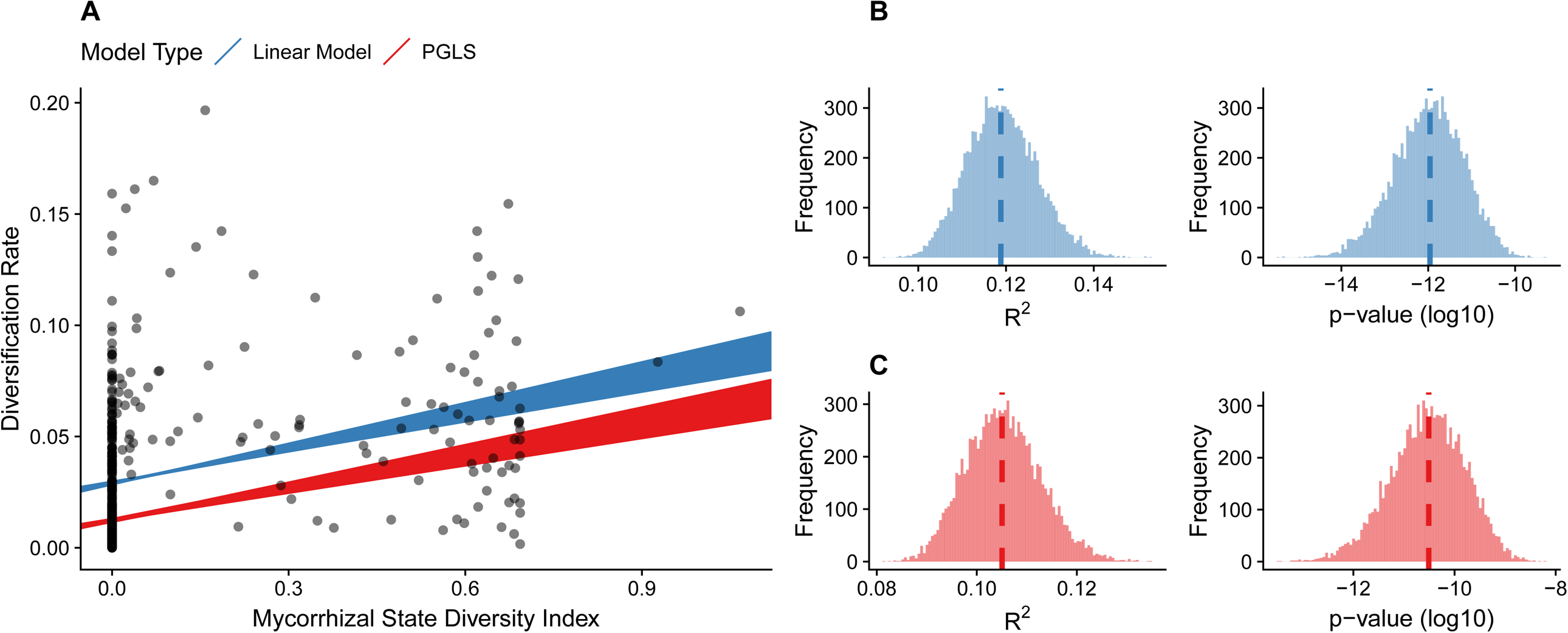
(a) Scatterplots showing an exemplar (1 out of 10000) relationship between mycorrhizal state diversity index and diversification rates estimated with □ (relative extinction fraction) = 0.9. The red and blue lines indicate the results of linear models and phylogenetic generalized least squares (PGLS) fit, respectively, for 10000 datasets of seed plant mycorrhizal states. Panels b and c show the frequency distribution of R^2^ and p-values of (b) linear models (blue bars) and (c) PGLS (red bars), dashed lines show the median for each distribution.

The additional analyses of adding a mycorrhizal misidentification to 20% of the species, supported our main conclusions, which are the positive association between mycorrhizal diversity index and diversification rates, and MIX families having higher diversification rates (Figs S3 and S4). These results are also observed when we use the phylogeny of Harris & Davis (2016; Appendix S4). The additional analyses using the species level dataset also showed a positive association between mycorrhizal diversity index and diversification rates (dataset excluding remarks: Figure S6 and Table S5). Although the ANOVA analysis at the species level showed a similar tendency for most comparisons, it did not show significant differences on diversification rates among different mycorrhizal states for all thresholds (dataset excluding species with remarks: Fig. S5, and Tables S1 and S2).

## Discussion

The association with mycorrhizal fungi has been indicated as a key acquisition in the evolution of plants, nevertheless its effect on plants diversification has not been evaluated before. Here we presented the first attempt to assess the relationship between mycorrhizal associations and diversification rates of plants. Due to the undersampling of seed plants phylogeny and mycorrhizal state database, we used a simple and conservative approach that allows us to tackle this question. Our results showed that there was no difference on diversification rates between AM, EM and NM families (Fig. 2; Table S13 and S14). This shows that families that acquired novel mycorrhizal associations (EM and NM) do not have higher diversification rates than families that retained the ancestral state (AM), contrary to what was expected in a scenario of key innovation in mycorrhizal associations as a mechanism of diversification. Thus, regarding our first question, the lineages that established derived mycorrhizal associations do not differ in their diversification rates from AM families. Contrary, our analyses showed that families with mixed mycorrhizal state have higher diversification rates than AM and NM families (Fig. 2, Table S14). It is interesting to notice that mycorrhizal diverse families have not only higher diversification when compared to low diverse families with the ancestral state, but also higher rates than families that have switched from the ancestral state to one novel mycorrhizal state. Accordingly, the correlation between age and richness of the families (shown in Fig. S1) showed that all MIX families fall in the upper part of distribution, meaning that this group exhibit higher diversification rates.

In addition, there was a positive and significant association between mycorrhizal state diversity index and diversification rates, which does not depend on our categorical criteria of mycorrhizal state assignment to families. These associations with diversification rates, are observed when correcting or not for the phylogenetic structure, suggesting that the relationship is not due to phylogenetic relatedness between families. Also, the patterns are observed under different scenarios of extinction, and even with □ = 0.0, where extinction is absent, the relationship is conserved. Given that diversification rates are determined by age and richness of the family, the effect of those variables could in theory have driven the relationship between mycorrhizal heterogeneity and diversification rates. We observed no significant correlation between mycorrhizal heterogeneity and age; and we see a similar pattern with species richness, and although the correlation is significant, the R^2^ is quite low (Fig. S2). This supports that mycorrhizal heterogeneity is mainly associated with diversification rates, not with age or richness per se. In addition, these patterns are also observed when we analyzed the species-level data (Appendix S5), which although has a lot less datapoints is supposedly more precise.

The mycorrhizal diversity index might not capture well the effects of mycorrhizal shifts on diversification rates if shifts occurred only once within each family. However, we observed that mycorrhizal shifts in mix families occurred multiple times, because more than 97% of mixed families contain genera that have multiple mycorrhizal states, which means that the different mycorrhizal states do not form monophyletic sub-clades and shifts occur even below the genus level. This suggests that diversification rates are not the result of a single mycorrhizal shift, but a result of high lability of the mycorrhizal states within mix families.

Both results, the ANOVA for family mycorrhizal state and association between mycorrhizal heterogeneity (MSDI) and diversification, suggest that independent of which mycorrhizal state is involved, a higher heterogeneity of mycorrhizal states in a family might promote diversification rates. These results together suggest that rather than a key innovation scenario, it is the evolutionary variability of mycorrhizal state what promotes diversification rates of plant seed families. We interpret mycorrhizal heterogeneity to result from a higher evolutionary lability of the mycorrhizal states within these families, which has been suggested to promote diversification in other biotic interactions (Hardy and Otto 2014). Different mycorrhizal state provides advantages to plants in certain environments but not in others (Brundrett, 2002), thus families that are composed by species with different mycorrhizal states might have been able to switch states in evolutionary time, making them able to evolve a higher diversity of niches which would result in a higher diversification rate. Under this scenario, mycorrhizal diverse families would have had more chances to take advantage of a new ecological opportunity, than families with most species within a single mycorrhizal state, resulting in broader range sizes, which has been demonstrated to explained most of variation in diversification rates among seed plant families (Hernandez & Wiens, 2020). Indeed, at the species level, facultatively mycorrhizal (AM-NM) species show the widest geographic and ecological amplitude (Hampel et al., 2013) and are more successful during naturalization than those with obligate mycorrhizal associations (Pysek et al 2019). Similarly, species that present facultative dual (EM-AM) associations have the higher percentage of naturalized species outside their native ranges (Moyano et al., 2020) and it has been suggested that the ability of species to form AM and EM associations may contribute to its wide geographic distribution, since they could use a wider range of habitats (Teste et al., 2019). The association between multiple mycorrhizal states and wider ranges observed at the species level is expected to occur also at genus and family level, since families containing species with different mycorrhizal states will be able to colonize a higher diversity of environmental conditions. Moreover, our results also highlight the evolutionary role of specialization at different organization levels: even if species are mycorrhizal specialized within a mixed family, the possibility to switch to different mycorrhizal states might increase the diversification of the family.

### Robustness of results

Since large database can potentially present some uncertainties, it is necessary to take them into account in order to test the robustness of the results. In this study we performed several sensitivity analyses to evaluate whether the results were maintained under different scenarios. In particular, we (1) generated 10000 datasets to properly address the uncertainty regarding the proportions of species in each mycorrhizal state for mixed genera; (2) introduced errors to the classification of mycorrhizal state on 20% of plant species (one order of magnitude higher than previous estimates of error for similar mycorrhizal databases); (3) tested different thresholds for mycorrhizal state assignation for each family; and (4) performed a similar analyses using the species-level database from the FungalRoot. Given that the observed patterns were maintained across all these tests, the sensitivity analysis suggest our results are extremely robust. We suggest that this approach to handle uncertainty can serve to the discussion of ways towards dealing with large datasets, in which one does not need to make arbitrary decisions but rather try different scenarios to investigate if the same patterns can be observed when taking into account different potential limitations of large datasets.

### Methodological caveats

Because biodiversity dynamics could be rather complex, with clades either expanding, at equilibrium and even declining in diversity, simple metrics like the average rate of diversification might not be able to separate them (Quental and Marshall 2010). The use of an average rate as a descriptor of a clade diversification dynamics assumes (or at least equates to) a scenario of expanding diversity (Quental and Marshall 2010), and it might be especially problematic if lineages have a carrying capacity because the average rate might be diluted as time goes by (Rabosky 2009). Moreover, with an average rate is not possible to distinguish between speciation and extinction rates or to test directly the effect of one trait on shifts in diversification rates. Ideally one would use more complex tests, but that would require a lot of more phylogenetic data than what is currently available. Overall, we used a relatively simple and limited macroevolutionary method that allowed us to compare diversification rates across clades with different mycorrhizal states and to evaluate the correlation between mycorrhizal heterogeneity and diversification, and our conclusions arose from a limited ecological and phylogenetic data. These conclusions might be revisited in future studies, when more data on mycorrhizal states and more complete phylogenies of plants are available.

## Conclusions

Acknowledging the limitations of our study, the results showed a positive relationship between mycorrhizal state diversity and diversification rates of seed plant families. This relationship was observed classifying families into a single mycorrhizal state (AM, EM, NM or MIX) and by calculating the heterogeneity of states using MSDI, both using the genus-level list and the species-level list from the Fungal Root database. The pattern is maintained including aleatory errors into both lists and testing different thresholds for family mycorrhizal state assignation. Finally, the results are also maintained when we performed same analyses using another seed plant phylogeny. The robustness of the results suggest that a higher diversity of mycorrhizal states promotes diversification of lineages, possibly related with new ecological opportunities that each mycorrhizal state provides to plants. Our results finally suggest that the associations between soil fungi and plants has been key for plant diversification, not only due to the foundational association that allows plants colonize land (Pirozynski and Malloch 1975) but also for further diversification of seed plant lineages.

## Acknowledgements

We thank all members of the Lab MeMe in the Institute of BioScience at the University of São Paulo for discussions and suggestions. TQ thanks FAPESP for financial support (grants # 2012/04072-3 and #2018/05462-6), GB thanks FAPESP for financial support (grant # 2018/04821-2) and MIM thanks CONICYT for the doctoral scholarship # 21151009.

## Data Accessibility

All data, R scripts, and documents related to this manuscript are available at [https://github.com/gburin/mycoDiversif]

