## Supplementary material for "Seed plant families with diverse mycorrhizal states have higher diversification rates"

#### Contents

**Appendix S1 - Comparison with background rates** **2**

**Appendix S2 - Relationship bewteen mycorrhizal diversity index, species richness and age** **3**

**Appendix S3 - Randomly adding 20% of misassignment of mycorrhizal type into the data obtained from genus-level list** **4**

**Appendix S4 - Analyses with the Species-level dataset** **5**

**Appendix S6 - Replicating main analyses with tree from Harris and Davies (2016)** **24**

**Appendix S7 - Phylogenetic signal of diversification rates, age, and richness** **26**

**Appendix S8 - Percentage of MIX species** **27**

#### Appendix S1 - Comparison with background rates

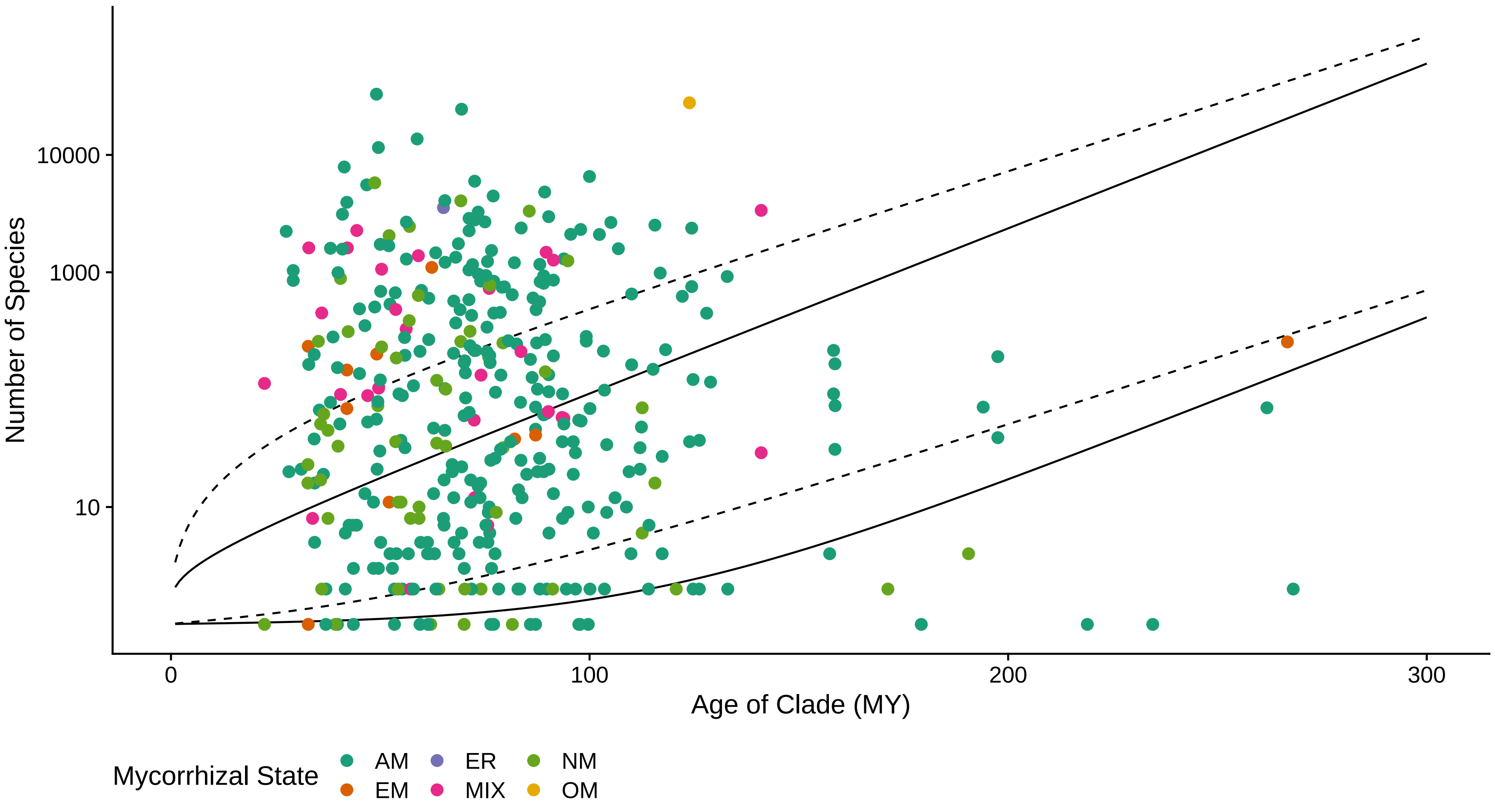

Figure S1: Relationship between number of species and age for each lineage when compared to the confidence intervals based on the global diversification rate of seed plants. The solid and dashed lines represent the expected richness for $\epsilon = 0$ and $\epsilon = 0.9$, respectively. The color of the points represents the mycorrhizal state for the 60% threshold.

### Appendix S2 - Relationship bewteen mycorrhizal diversity index, species richness and age

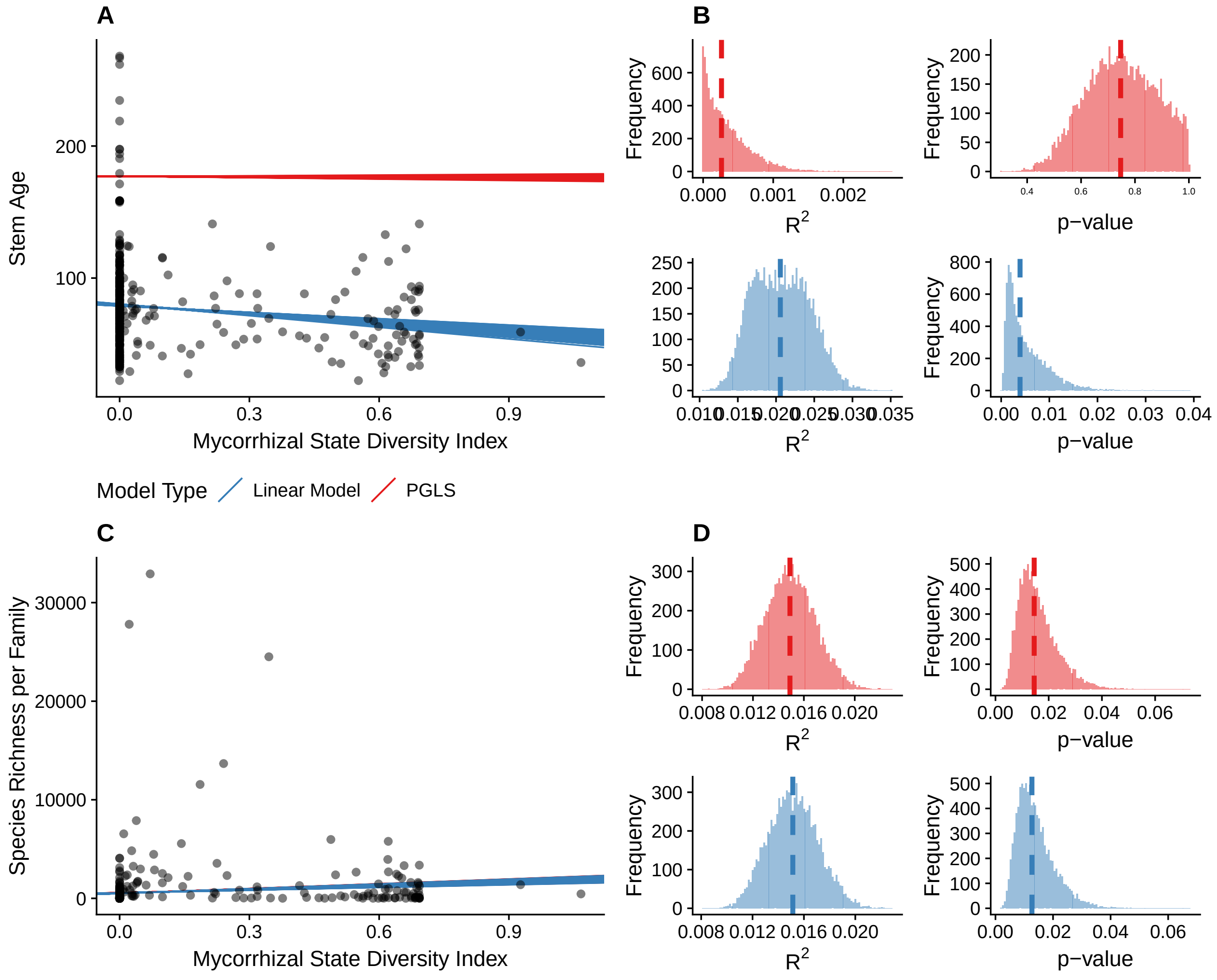

Figure S2: Scatterplots showing the relationship between mycorrhizal diversity index and (a) stem age and (c) species richness. Panels (b) and (d) show the frequency distribution of $R^2$ and p-values of linear models (blue bars) and PGLS (red bars). Dashed lines show the median for each distribution.

### Appendix S3 - Randomly adding 20% of misassignment of mycorrhizal type into the data obtained from genus-level list

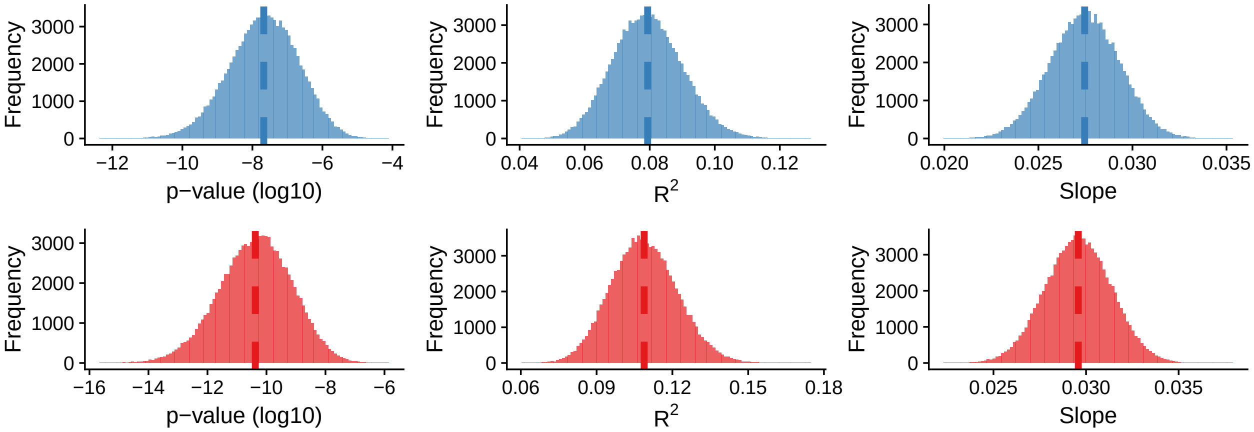

Figure S3: Distribution of $R^2$, p-value and slope value distributions for $\epsilon = 0$ after adding 20% misassignment of mycorrhizal type. Blue histograms (upper row) indicate results for standard linear models, whereas red histograms (lower row) indicate results for phylogenetic generalized least squares.

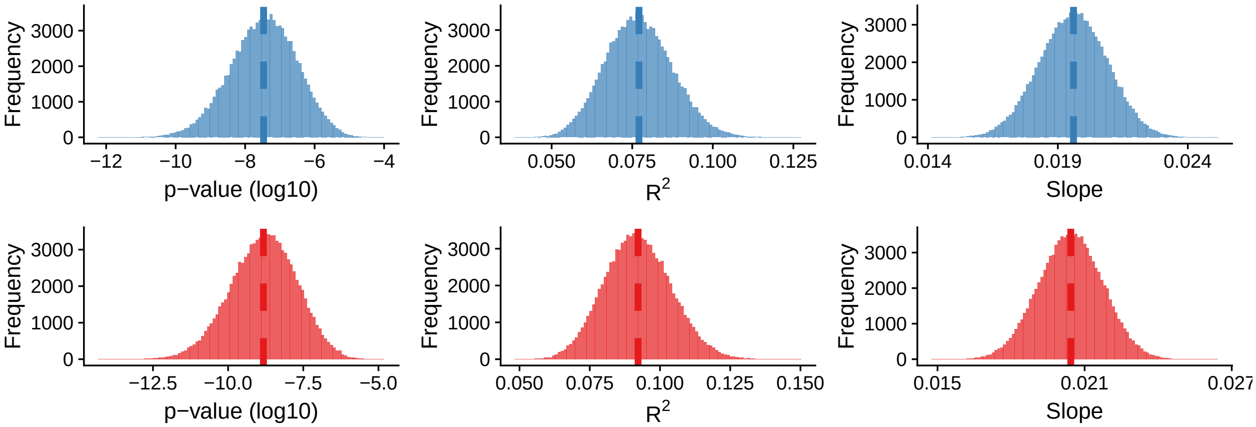

Figure S4: Distribution of $R^2$, p-value and slope value distributions for $\epsilon = 0.9$ after adding 20% misassignment of mycorrhizal type. Blue histograms (upper row) indicate results for standard linear models, whereas red histograms (lower row) indicate results for phylogenetic generalized least squares.

### Appendix S4 - Analyses with the Species-level dataset

#### Clean dataset - excluding species with any remark

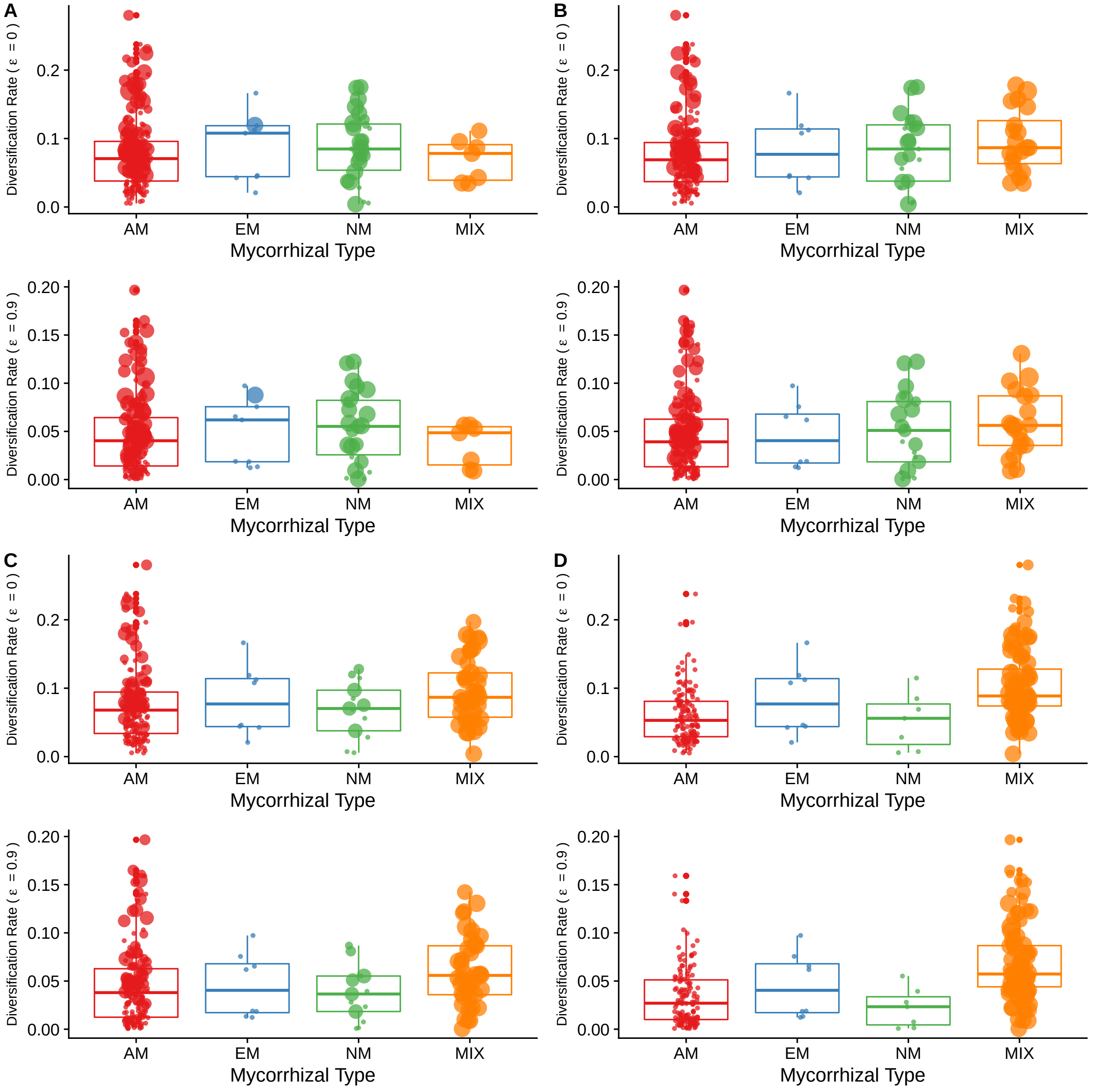

Figure S5: Relationship between mycorrhizal type and diversification rates using the thresholds (a) 50%, (b) 60%, (c) 80% and (d) 100% for MIX state assignment using the species-level dataset after removing species with remarks. Each panel shows diversification rate estimated with $\epsilon = 0$ (upper) and diversification rate estimated with $\epsilon = 0.9$ (lower). AM: Arbuscular mycorrhiza, EM: Ectomycorrhiza, NM: non-mycorrhizal and MIX (families with no dominance of any specific mycorrhizal association). The size of the points indicates the Mycorrhizal Type Diversity Index value for each lineage, indicating a predominance of larger indices with higher diversification rates.

##### Summary statistics using clean species-level dataset

Table S1: Summary statistics for phyANOVA for both values of $\epsilon$ using the species-level dataset after removing species with remarks to test for significant differences in diversification rates. Significant values are highlighted in bold.

| | e = 0 | | e = 0.9 | |
|---|---|---|---|---|
| Threshold | F | p-value | F | p-value |
| 50% | 0.385 | 0.864 | 0.494 | 0.811 |
| 60% | 1.538 | 0.394 | 0.928 | 0.625 |
| 80% | 3.784 | 0.093 | 3.262 | 0.122 |
| 100% | 24.583 | **0.001** | 27.097 | **0.001** |

Table S2: Summary statistics for standard ANOVA for both values of $\epsilon$ using the species-level dataset after removing species with remarks to test for significant differences in diversification rates. Significant values are highlighted in bold.

| | e = 0 | | e = 0.9 | |
|---|---|---|---|---|
| Threshold | F | p-value | F | p-value |
| 50% | 0.396 | 0.756 | 0.501 | 0.682 |
| 60% | 1.563 | 0.199 | 0.948 | 0.418 |
| 80% | 3.529 | **0.015** | 2.997 | **0.031** |
| 100% | 23.996 | **0** | 26.323 | **0** |

##### Posthoc tests using clean species-level dataset

Table S3: Pairwise Corrected p-values for phyANOVA using the species-level dataset after removing species with remarks. Significant values are highlighted in bold.

| | | e = 0 | | | | e = 0.9 | | | |
|---|---|---|---|---|---|---|---|---|---|
| Threshold | Mycorrhizal Type | AM | EM | MIX | NM | AM | EM | MIX | NM |
| 50% | AM | 1 | 1 | 1 | 1 | 1 | 1 | 1 | 1 |
| 50% | EM | 1 | 1 | 1 | 1 | 1 | 1 | 1 | 1 |
| 50% | MIX | 1 | 1 | 1 | 1 | 1 | 1 | 1 | 1 |
| 50% | NM | 1 | 1 | 1 | 1 | 1 | 1 | 1 | 1 |
| 60% | AM | 1 | 1 | 0.24 | 1 | 1 | 1 | 0.732 | 1 |
| 60% | EM | 1 | 1 | 1 | 1 | 1 | 1 | 1 | 1 |
| 60% | MIX | 0.24 | 1 | 1 | 1 | 0.732 | 1 | 1 | 1 |
| 60% | NM | 1 | 1 | 1 | 1 | 1 | 1 | 1 | 1 |
| 80% | AM | 1 | 1 | **0.042** | 1 | 1 | 1 | 0.12 | 1 |
| 80% | EM | 1 | 1 | 1 | 1 | 1 | 1 | 1 | 1 |
| 80% | MIX | **0.042** | 1 | 1 | 0.12 | 0.12 | 1 | 1 | 0.12 |
| 80% | NM | 1 | 1 | 0.12 | 1 | 1 | 1 | 0.12 | 1 |
| 100% | AM | 1 | 0.777 | **0.006** | 0.777 | 1 | 0.72 | **0.006** | 0.72 |
| 100% | EM | 0.777 | 1 | 0.76 | 0.777 | 0.72 | 1 | 0.356 | 0.72 |
| 100% | MIX | **0.006** | 0.76 | 1 | **0.006** | **0.006** | 0.356 | 1 | **0.006** |
| 100% | NM | 0.777 | 0.777 | **0.006** | 1 | 0.72 | 0.72 | **0.006** | 1 |

Table S4: Pairwise Corrected p-values for standard ANOVA. Significant values are highlighted in bold.

| | e = 0 | | | | e = 0.9 | | | |
|---|---|---|---|---|---|---|---|---|
| Types | 50% | 60% | 80% | 100% | 50% | 60% | 80% | 100% |
| EM-AM | 0.964 | 0.992 | 0.985 | 0.613 | 0.993 | 1 | 1 | 0.78 |
| MIX-AM | 0.995 | 0.146 | **0.012** | **0** | 0.858 | 0.348 | **0.036** | **0** |
| NM-AM | 0.776 | 0.938 | 0.911 | 0.808 | 0.831 | 0.974 | 0.778 | 0.626 |
| MIX-EM | 0.954 | 0.769 | 0.704 | 0.237 | 0.866 | 0.768 | 0.622 | 0.106 |
| NM-EM | 1 | 1 | 0.897 | 0.411 | 0.996 | 0.993 | 0.927 | 0.404 |
| NM-MIX | 0.892 | 0.632 | 0.09 | **0** | 0.663 | 0.78 | 0.081 | **0** |

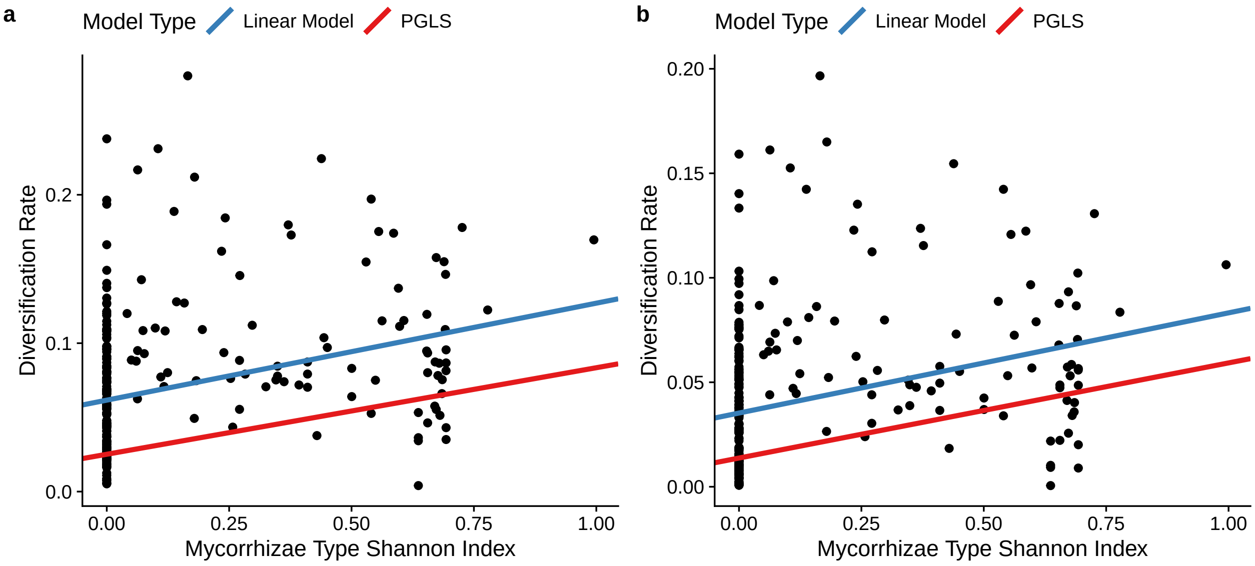

Figure S6: Scatterplots showing the relationship between mycorrhizal diversity index and diversification rates using the species-level dataset after removing species with remarks. Diversification rates were estimated with $\epsilon = 0$ (a) and with $\epsilon = 0.9$ (b). The red and blue lines indicate the results of a linear model and a phylogenetic generalized least squares (PGLS) fit, respectively.

Table S5: Summary statistics for the phylogenetic and parametric regressions using the species-level dataset after removing species with remarks. Significant values are highlighted in bold.

| | e = 0 | | e = 0.9 | |
|---|---|---|---|---|
| Model | p-value | $R^2$ | p-value | $R^2$ |
| PGLS | **4.063e-07** | 0.08904 | **3.483e-07** | 0.09007 |
| LM | **1.676e-07** | 0.09458 | **2.844e-07** | 0.09109 |

#### Full species-level dataset - including species with any remark

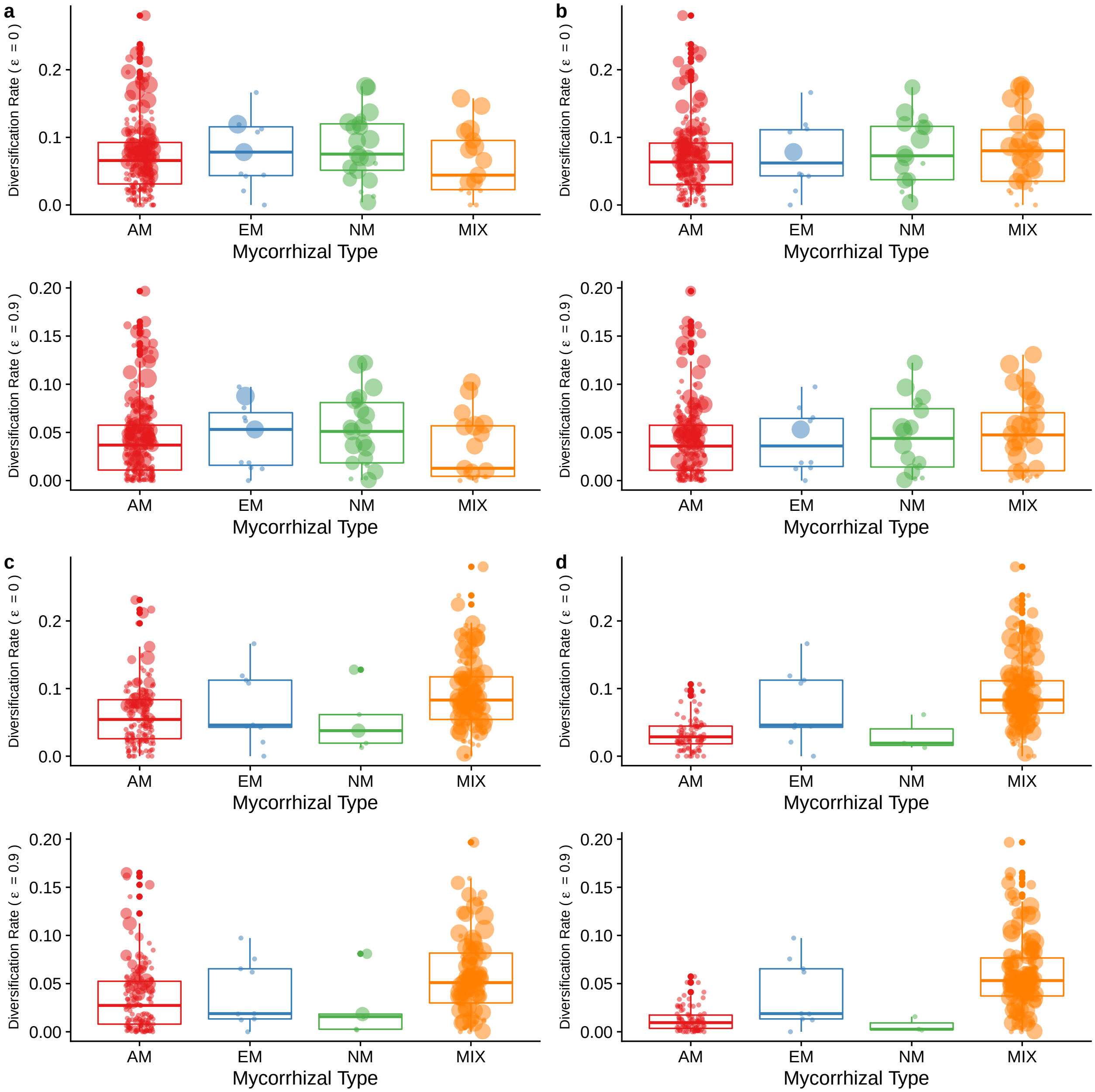

Figure S7: Relationship between mycorrhizal type and diversification rates using the thresholds (a) 50%, (b) 60%, (c) 80% and (d) 100% for MIX state assignment using the species-level dataset with species with remarks. Each panel shows diversification rate estimated with $\epsilon = 0$ (upper) and diversification rate estimated with $\epsilon = 0.9$ (lower). AM: Arbuscular mycorrhiza, EM: Ectomycorrhiza, NM: non-mycorrhizal and MIX (families with no dominance of any specific mycorrhizal association). The size of the points indicates the Mycorrhizal Type Diversity Index value for each lineage, indicating a predominance of larger indices with higher diversification rates.

##### Summary statistics using full species-level dataset

Table S6: Summary statistics for phyANOVA for both values of $\epsilon$ using the species-level dataset with species with remarks to test for differences in diversification rates. Significant values are highlighted in bold.

| | e = 0 | | e = 0.9 | |
|---|---|---|---|---|
| Threshold | F | p-value | F | p-value |
| 50 | 0.691 | 0.655 | 0.666 | 0.699 |
| 60 | 0.511 | 0.795 | 0.324 | 0.877 |
| 80 | 8.571 | **0.003** | 8.425 | **0.001** |
| 100 | 36.44 | **0.001** | 43.806 | **0.001** |

Table S7: Summary statistics for standard ANOVA for both values of $\epsilon$ using the species-level dataset with species with remarks to test for differences in diversification rates. Significant values are highlighted in bold.

| | e = 0 | | e = 0.9 | |
|---|---|---|---|---|
| Threshold | F | p-value | F | p-value |
| 50 | 1.083 | 0.365 | 1.085 | 0.364 |
| 60 | 0.953 | 0.434 | 0.831 | 0.506 |
| 80 | 7.113 | **0** | 7.084 | **0** |
| 100 | 28.22 | **0** | 34.224 | **0** |

##### Posthoc tests using full species-level dataset

Table S8: Pairwise Corrected p-values for phyANOVA using the species-level dataset with species with remarks. Significant values are highlighted in bold.

| | | e = 0 | | | | e = 0.9 | | | |
|---|---|---|---|---|---|---|---|---|---|
| Threshold | Mycorrhizal Type | AM | EM | MIX | NM | AM | EM | MIX | NM |
| 50% | AM | 1 | 1 | 1 | 1 | 1 | 1 | 1 | 1 |
| 50% | EM | 1 | 1 | 1 | 1 | 1 | 1 | 1 | 1 |
| 50% | MIX | 1 | 1 | 1 | 1 | 1 | 1 | 1 | 0.81 |
| 50% | NM | 1 | 1 | 1 | 1 | 1 | 1 | 0.81 | 1 |
| 60% | AM | 1 | 1 | 1 | 1 | 1 | 1 | 1 | 1 |
| 60% | EM | 1 | 1 | 1 | 1 | 1 | 1 | 1 | 1 |
| 60% | MIX | 1 | 1 | 1 | 1 | 1 | 1 | 1 | 1 |
| 60% | NM | 1 | 1 | 1 | 1 | 1 | 1 | 1 | 1 |
| 80% | AM | 1 | 1 | **0.006** | 1 | 1 | 1 | **0.006** | 1 |
| 80% | EM | 1 | 1 | 1 | 1 | 1 | 1 | 1 | 1 |
| 80% | MIX | **0.006** | 1 | 1 | 0.31 | **0.006** | 1 | 1 | 0.185 |
| 80% | NM | 1 | 1 | 0.31 | 1 | 1 | 1 | 0.185 | 1 |
| 100% | AM | 1 | 0.184 | **0.006** | 0.93 | 1 | 0.224 | **0.006** | 0.778 |
| 100% | EM | 0.184 | 1 | 0.664 | 0.525 | 0.224 | 1 | 0.405 | 0.405 |
| 100% | MIX | **0.006** | 0.664 | 1 | **0.045** | **0.006** | 0.405 | 1 | **0.015** |
| 100% | NM | 0.93 | 0.525 | **0.045** | 1 | 0.778 | 0.405 | **0.015** | 1 |

Table S9: Pairwise Corrected p-values for standard ANOVA. Significant values are highlighted in bold.

| | e = 0 | | | | e = 0.9 | | | |
|---|---|---|---|---|---|---|---|---|
| Types | 50% | 60% | 80% | 100% | 50% | 60% | 80% | 100% |
| EM-AM | 0.991 | 0.999 | 0.937 | 0.067 | 0.998 | 1 | 0.99 | 0.084 |
| ER-AM | 0.592 | 0.62 | 0.808 | 1 | 0.562 | 0.59 | 0.783 | 1 |
| MIX-AM | 0.97 | 0.805 | **0** | **0** | 0.906 | 0.874 | **0** | **0** |
| NM-AM | 0.763 | 0.958 | 0.996 | 1 | 0.85 | 0.995 | 0.968 | 0.998 |
| ER-EM | 0.561 | 0.663 | 0.649 | 0.527 | 0.593 | 0.723 | 0.735 | 0.601 |
| MIX-EM | 0.937 | 0.996 | 0.778 | 0.674 | 0.929 | 0.986 | 0.608 | 0.364 |
| NM-EM | 0.997 | 0.999 | 0.935 | 0.583 | 0.996 | 0.999 | 0.928 | 0.485 |
| MIX-ER | 0.838 | 0.407 | 0.12 | 0.05 | 0.87 | 0.413 | 0.113 | **0.029** |
| NM-ER | 0.352 | 0.501 | 0.979 | 1 | 0.368 | 0.561 | 0.993 | 0.999 |
| NM-MIX | 0.684 | 1 | 0.358 | 0.102 | 0.637 | 0.999 | 0.237 | **0.03** |

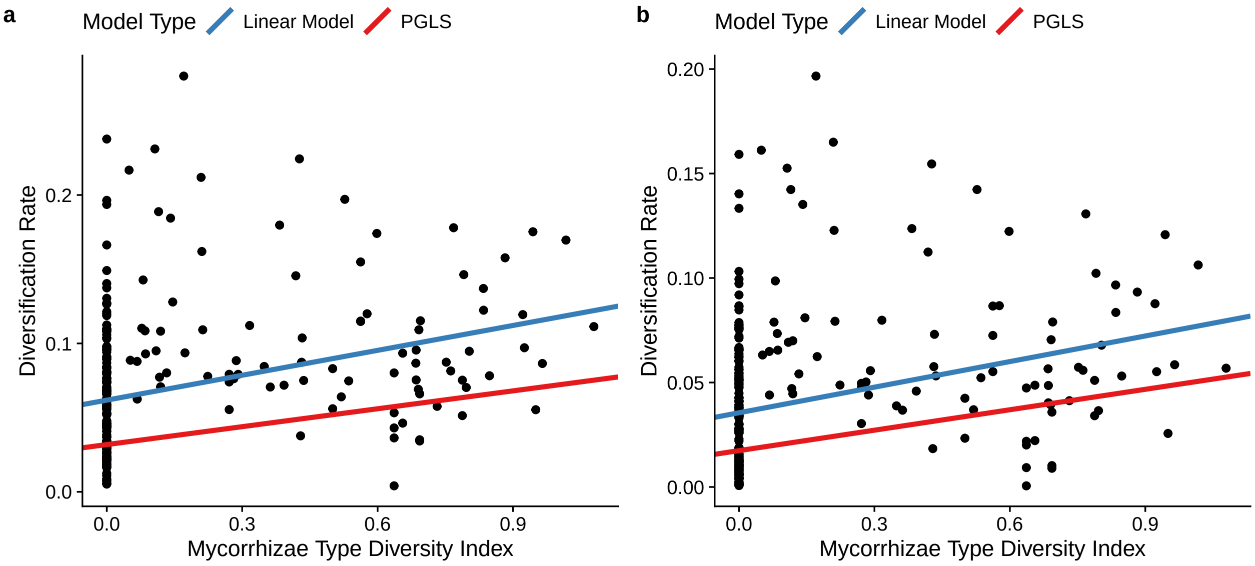

Figure S8: Scatterplots showing the relationship between mycorrhizal diversity index and diversification rates using the species-level dataset with species with remarks. Diversification rates were estimated with $\epsilon = 0$ (a) and with $\epsilon = 0.9$ (b). The red and blue lines indicate the results of a linear model and a phylogenetic generalized least squares (PGLS) fit, respectively.

Table S10: Summary statistics for the phylogenetic and parametric regressions using the species-level dataset with species with remarks. Significant values are highlighted in bold.

| | e = 0 | | e = 0.9 | |
|---|---|---|---|---|
| Model | p-value | $R^2$ | p-value | $R^2$ |
| PGLS | **5.265e-05** | 0.06444 | **4.013e-05** | 0.06653 |
| LM | **2.528e-07** | 0.09354 | **4.977e-07** | 0.08898 |

### Appendix S5 - Analyses with the genus-level dataset

#### Thresholds for mycorrhizal state assignment

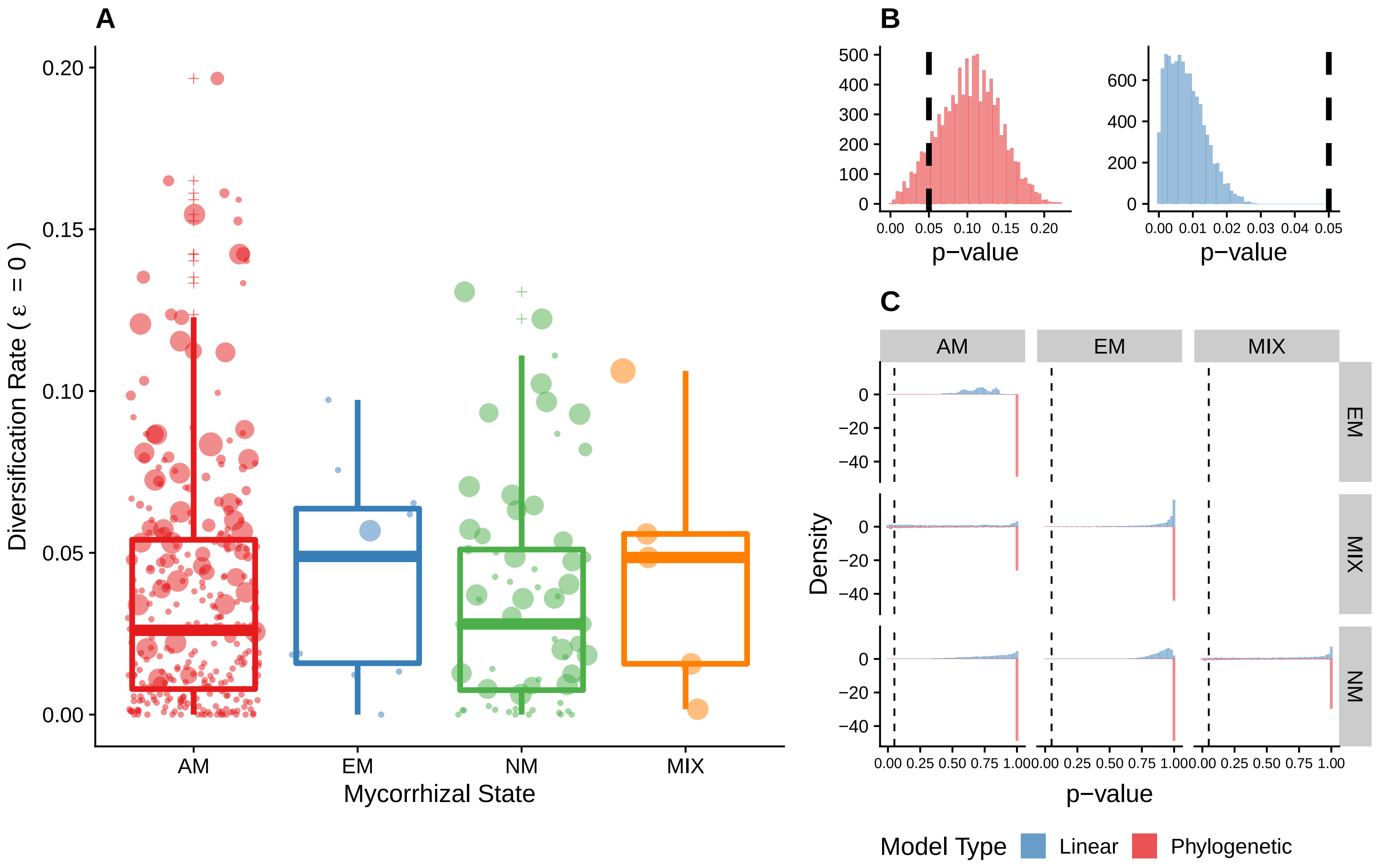

Figure S9: Relationship between mycorrhizal type and diversification rates using the threshold 50% for MIX state assignment using the genus-level dataset and with $\epsilon = 0$. Panel (A): Boxplot with a randomly chosen dataset replicate showing the distribution of diversification rates for each mycorrhizal state. AM: Arbuscular mycorrhiza, EM: Ectomycorrhiza, NM: non-mycorrhizal and MIX (families with no dominance of any specific mycorrhizal association). The size of the points indicates the Mycorrhizal Type Diversity Index value for each lineage, indicating a predominance of larger indices with higher diversification rates. Panel (B): Histogram of p-value both phylogenetic (red) and standard (blue) ANOVA for all 10000 datasets. Panel (C): p-value of post-hoc tests of all pairwise combinations of mycorrhizal states using all 10000 datasets. In panels (B) and (C) the vertical dashed line correspond to 0.05

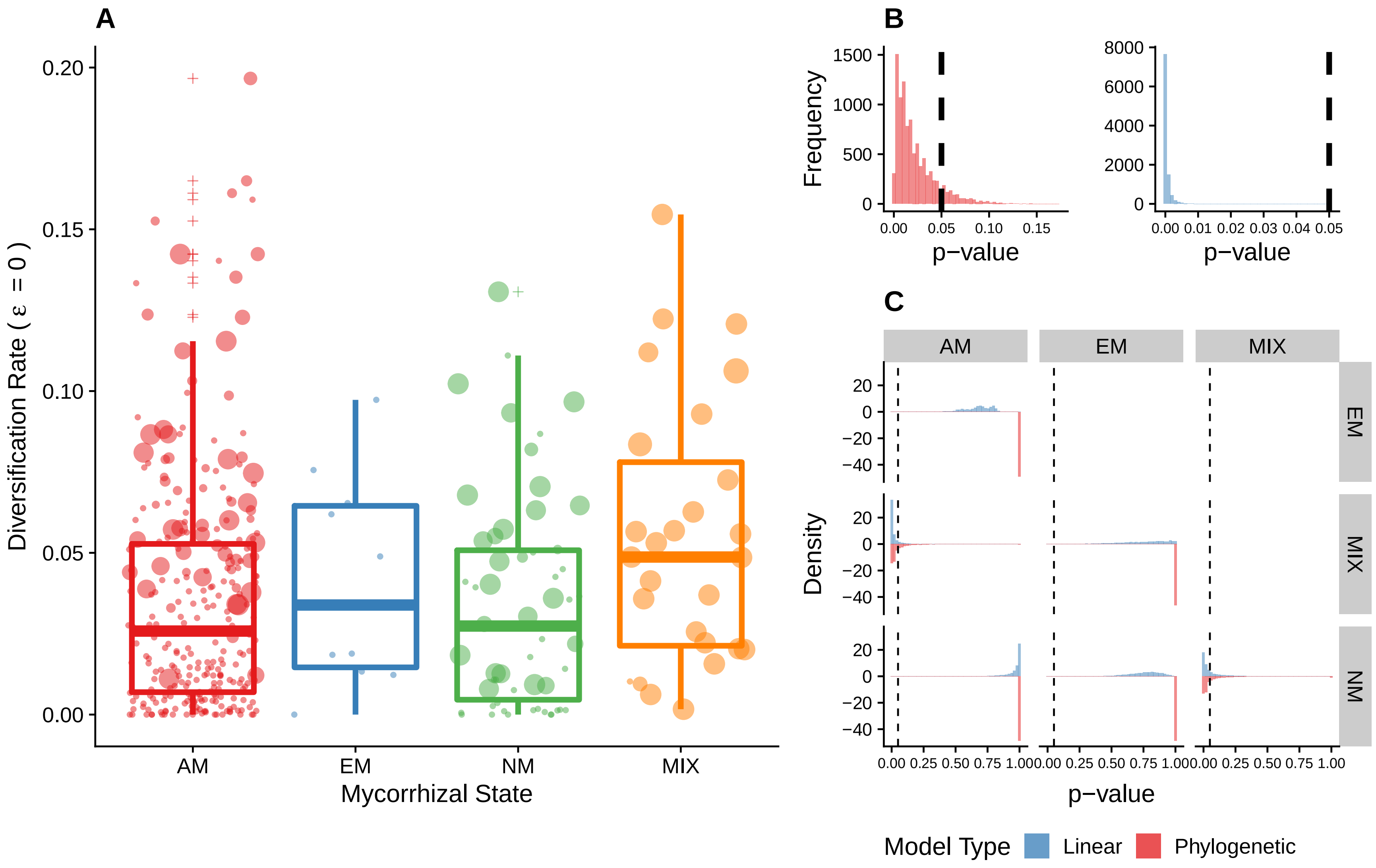

Figure S10: Relationship between mycorrhizal type and diversification rates using the threshold 60% for MIX state assignment using the genus-level dataset and with $\epsilon = 0$. Panel (A): Boxplot of randomly chosen dataset replicate showing the distribution of diversification rates for each mycorrhizal state. AM: Arbuscular mycorrhiza, EM: Ectomycorrhiza, NM: non-mycorrhizal and MIX (families with no dominance of any specific mycorrhizal association). The size of the points indicates the Mycorrhizal Type Diversity Index value for each lineage, indicating a predominance of larger indices with higher diversification rates. Panel (B): Histogram of p-value both phylogenetic (red) and standard (blue) ANOVA for all 10000 datasets. Panel (C): p-value of post-hoc tests of all pairwise combinations of mycorrhizal states using all 10000 datasets. In panels (B) and (C) the vertical dashed line correspond to 0.05

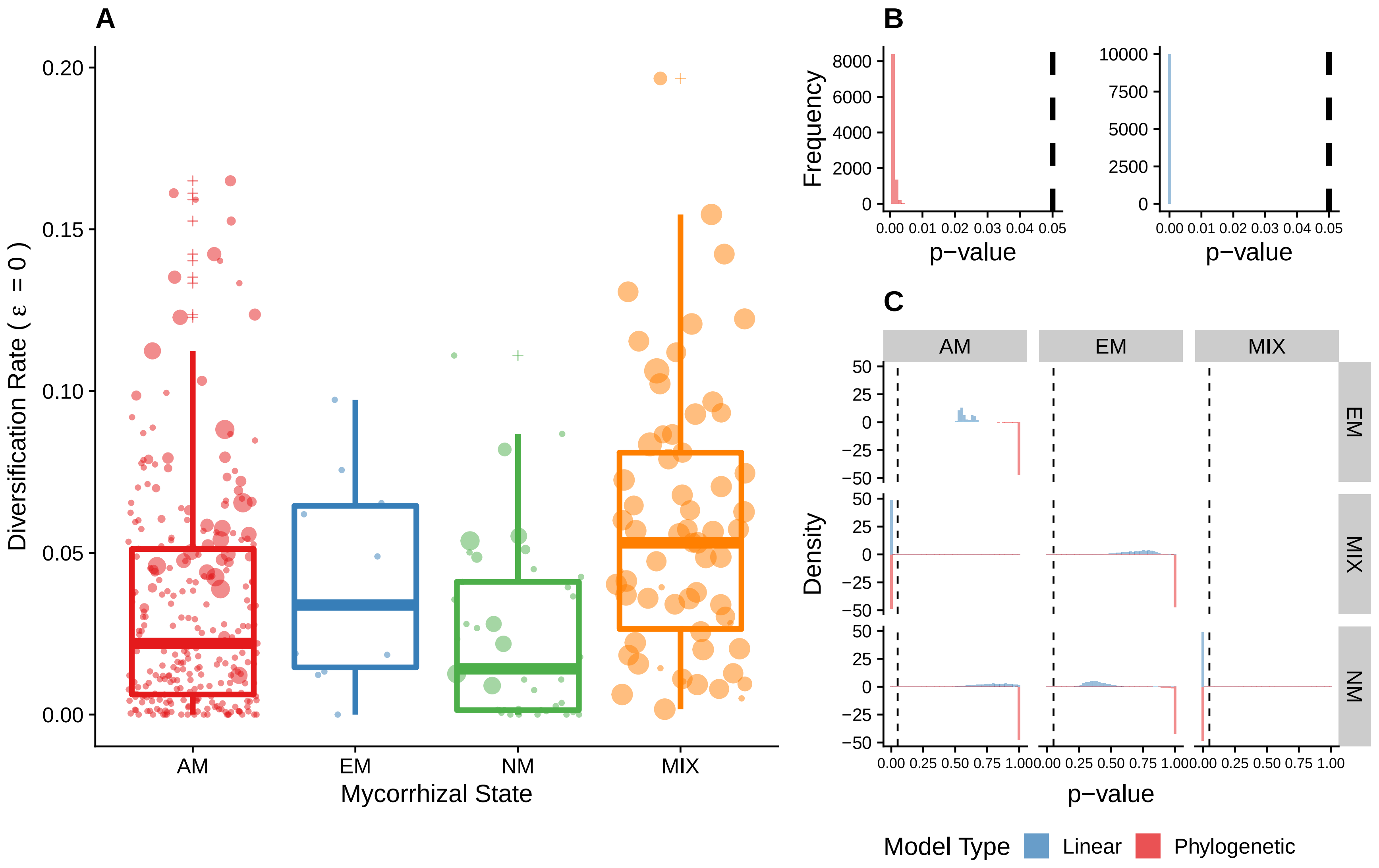

Figure S11: Relationship between mycorrhizal type and diversification rates using the threshold 80% for MIX state assignment using the genus-level dataset and with $\epsilon = 0$. Panel (A): Boxplot of randomly chosen dataset replicate showing the distribution of diversification rates for each mycorrhizal state. AM: Arbuscular mycorrhiza, EM: Ectomycorrhiza, NM: non-mycorrhizal and MIX (families with no dominance of any specific mycorrhizal association). The size of the points indicates the Mycorrhizal Type Diversity Index value for each lineage, indicating a predominance of larger indices with higher diversification rates. Panel (B): Histogram of p-value both phylogenetic (red) and standard (blue) ANOVA for all 10000 datasets. Panel (C): p-value of post-hoc tests of all pairwise combinations of mycorrhizal states using all 10000 datasets. In panels (B) and (C) the vertical dashed line correspond to 0.05

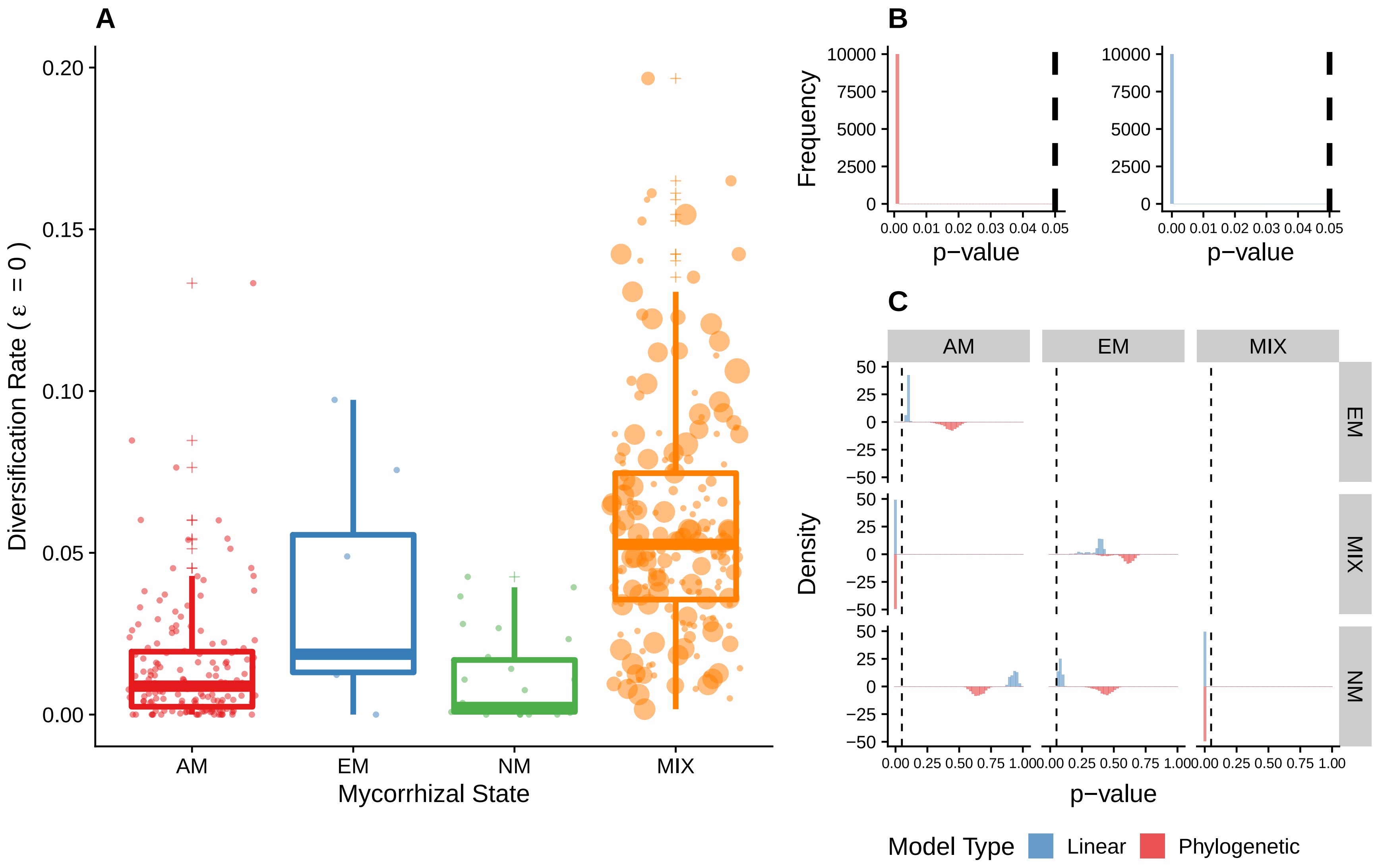

Figure S12: Relationship between mycorrhizal type and diversification rates using the threshold 100% for MIX state assignment using the genus-level dataset and with $\epsilon = 0$. Panel (A): Boxplot of randomly chosen dataset replicate showing the distribution of diversification rates for each mycorrhizal state. AM: Arbuscular mycorrhiza, EM: Ectomycorrhiza, NM: non-mycorrhizal and MIX (families with no dominance of any specific mycorrhizal association). The size of the points indicates the Mycorrhizal Type Diversity Index value for each lineage, indicating a predominance of larger indices with higher diversification rates. Panel (B): Histogram of p-value both phylogenetic (red) and standard (blue) ANOVA for all 10000 datasets. Panel (C): p-value of post-hoc tests of all pairwise combinations of mycorrhizal states using all 10000 datasets. In panels (B) and (C) the vertical dashed line correspond to 0.05

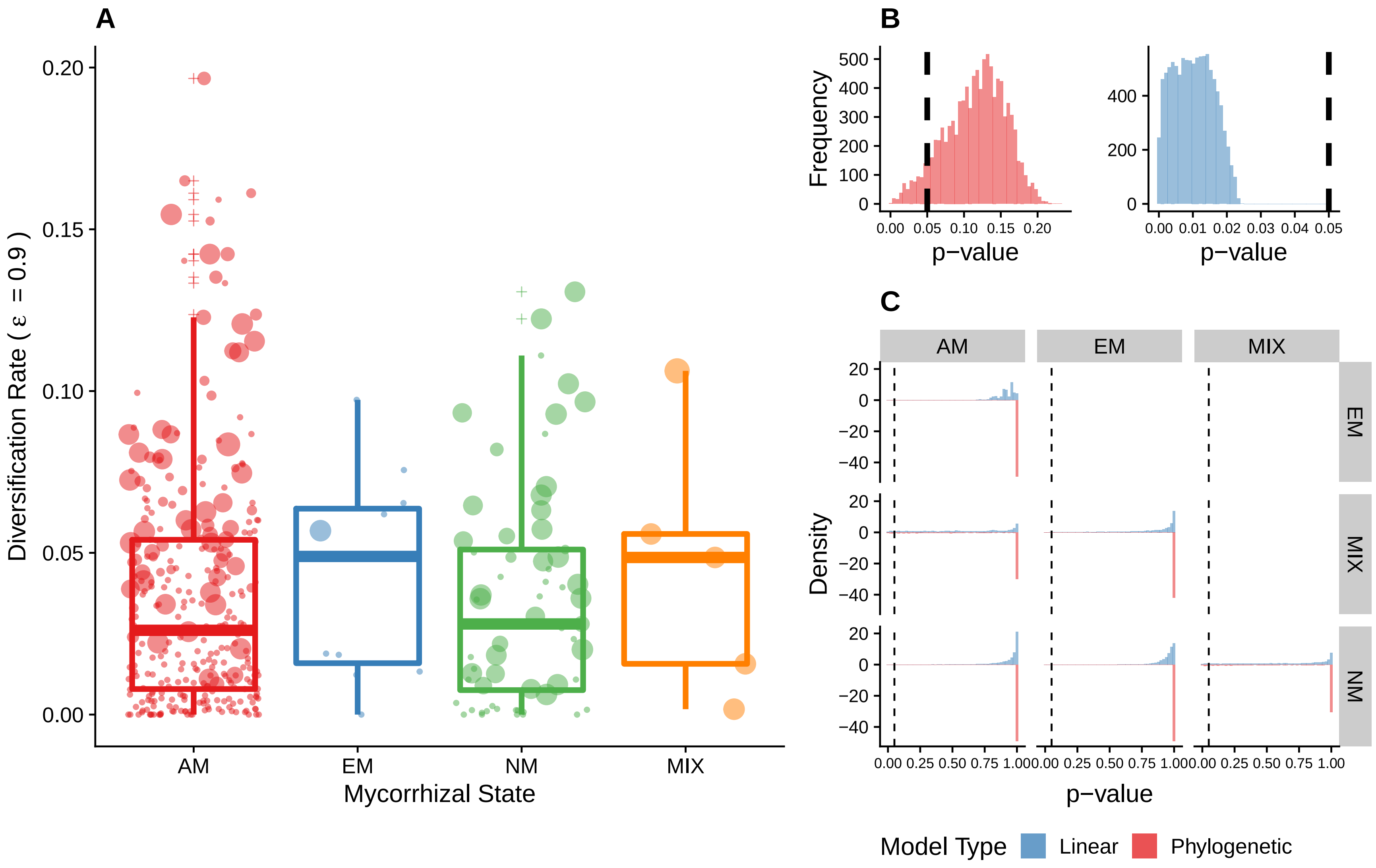

Figure S13: Relationship between mycorrhizal type and diversification rates using the threshold 50% for MIX state assignment using the genus-level dataset and with $\epsilon = 0.9$. Panel (A): Boxplot of randomly chosen dataset replicate showing the distribution of diversification rates for each mycorrhizal state. AM: Arbuscular mycorrhiza, EM: Ectomycorrhiza, NM: non-mycorrhizal and MIX (families with no dominance of any specific mycorrhizal association). The size of the points indicates the Mycorrhizal Type Diversity Index value for each lineage, indicating a predominance of larger indices with higher diversification rates. Panel (B): Histogram of p-value both phylogenetic (red) and standard (blue) ANOVA for all 10000 datasets. Panel (C): p-value of post-hoc tests of all pairwise combinations of mycorrhizal states using all 10000 datasets. In panels (B) and (C) the vertical dashed line correspond to 0.05

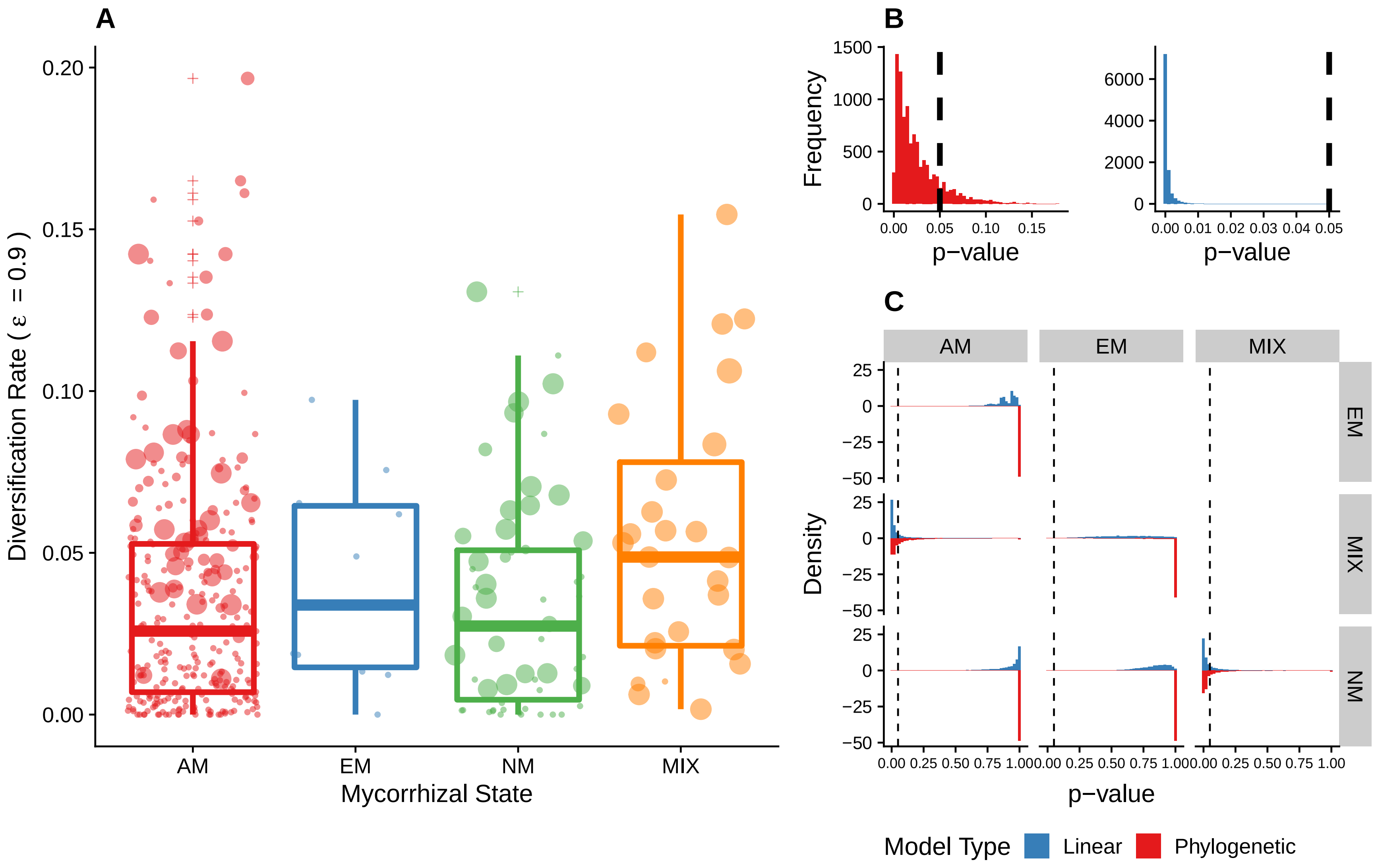

Figure S14: Relationship between mycorrhizal type and diversification rates using the threshold 60% for MIX state assignment using the genus-level dataset and with $\epsilon = 0.9$. Panel (A): Boxplot of randomly chosen dataset replicate showing the distribution of diversification rates for each mycorrhizal state. AM: Arbuscular mycorrhiza, EM: Ectomycorrhiza, NM: non-mycorrhizal and MIX (families with no dominance of any specific mycorrhizal association). The size of the points indicates the Mycorrhizal Type Diversity Index value for each lineage, indicating a predominance of larger indices with higher diversification rates. Panel (B): Histogram of p-value both phylogenetic (red) and standard (blue) ANOVA for all 10000 datasets. Panel (C): p-value of post-hoc tests of all pairwise combinations of mycorrhizal states using all 10000 datasets. In panels (B) and (C) the vertical dashed line correspond to 0.05. This is the same as figure 2 of the main text and was left there only for easy comparison.

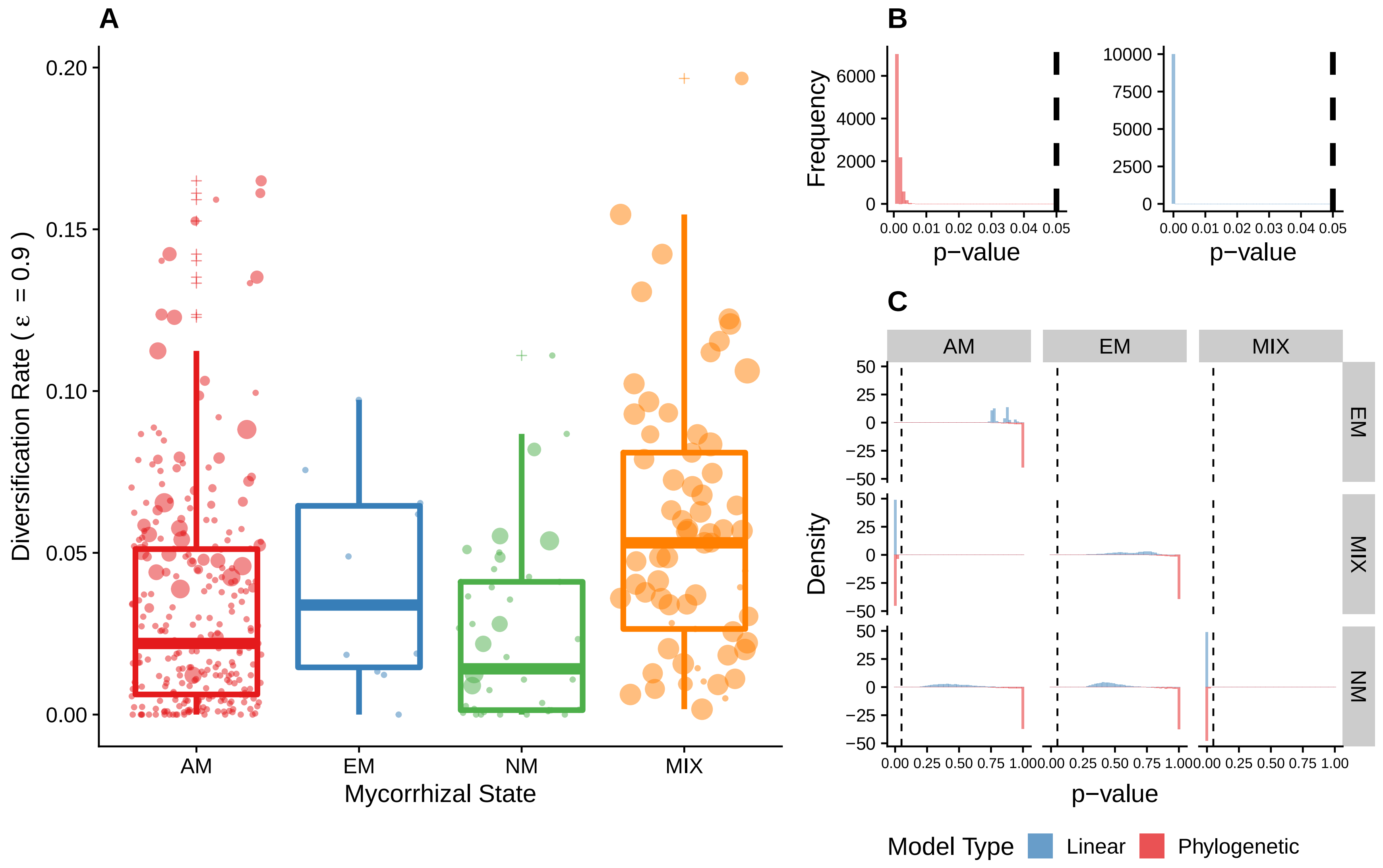

Figure S15: Relationship between mycorrhizal type and diversification rates using the threshold 80% for MIX state assignment using the genus-level dataset and with $\epsilon = 0.9$. Panel (A): Boxplot of randomly chosen dataset replicate showing the distribution of diversification rates for each mycorrhizal state. AM: Arbuscular mycorrhiza, EM: Ectomycorrhiza, NM: non-mycorrhizal and MIX (families with no dominance of any specific mycorrhizal association). The size of the points indicates the Mycorrhizal Type Diversity Index value for each lineage, indicating a predominance of larger indices with higher diversification rates. Panel (B): Histogram of p-value both phylogenetic (red) and standard (blue) ANOVA for all 10000 datasets. Panel (C): p-value of post-hoc tests of all pairwise combinations of mycorrhizal states using all 10000 datasets. In panels (B) and (C) the vertical dashed line correspond to 0.05

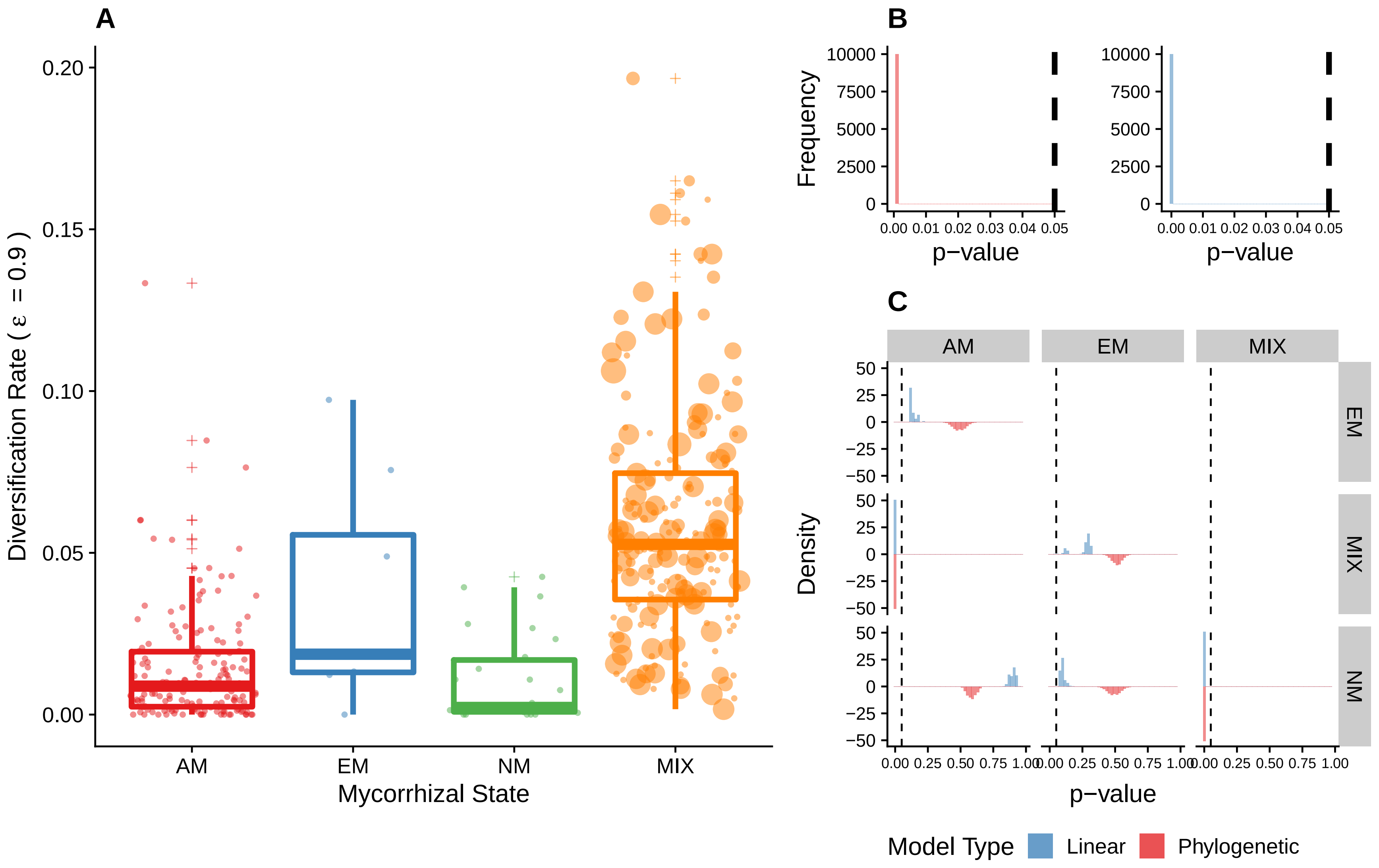

Figure S16: Relationship between mycorrhizal type and diversification rates using the threshold 100% for MIX state assignment using the genus-level dataset and with $\epsilon = 0.9$. Panel (A): Boxplot of randomly chosen dataset replicate showing the distribution of diversification rates for each mycorrhizal state. AM: Arbuscular mycorrhiza, EM: Ectomycorrhiza, NM: non-mycorrhizal and MIX (families with no dominance of any specific mycorrhizal association). The size of the points indicates the Mycorrhizal Type Diversity Index value for each lineage, indicating a predominance of larger indices with higher diversification rates. Panel (B): Histogram of p-value both phylogenetic (red) and standard (blue) ANOVA for all 10000 datasets. Panel (C): p-value of post-hoc tests of all pairwise combinations of mycorrhizal states using all 10000 datasets. In panels (B) and (C) the vertical dashed line correspond to 0.05

**Summary statistics**

Table S11: Summary statistics for phyANOVA for both values of $\epsilon$ using the genus-level dataset including all families to test for differences in diversification rates. Significant values are highlighted in bold.

| | e = 0 | | | | e = 0.9 | | | |
|---|---|---|---|---|---|---|---|---|
| Quantile | 50% | 60% | 80% | 100% | 50% | 60% | 80% | 100% |
| Min | **0.001** | **0.001** | **0.001** | **0.001** | **0.001** | **0.001** | **0.001** | **0.001** |
| 1st Quart | 0.075 | **0.007** | **0.001** | **0.001** | 0.088 | **0.008** | **0.001** | **0.001** |
| Median | 0.104 | **0.016** | **0.001** | **0.001** | 0.12 | **0.018** | **0.001** | **0.001** |
| Mean | 0.1026505 | **0.0228431** | **0.0011916** | **0.001** | 0.1155908 | **0.025796** | **0.0014098** | **0.001** |
| 3rd Quart | 0.131 | **0.032** | **0.001** | **0.001** | 0.146 | **0.036** | **0.002** | **0.001** |
| Max | 0.222 | 0.173 | **0.006** | **0.001** | 0.232 | 0.179 | **0.008** | **0.001** |

Table S12: Summary statistics for standard ANOVA for both values of $\epsilon$ using the genus-level dataset including all families to test for differences in diversification rates. Values were rounded to the fifth decimal place, and significant values are highlighted in bold.

| | e = 0 | | | | e = 0.9 | | | |
|---|---|---|---|---|---|---|---|---|
| Quantile | 50% | 60% | 80% | 100% | 50% | 60% | 80% | 100% |
| Minimum | **0** | **0** | **0** | **0** | **0** | **0** | **0** | **0** |
| 1st Quart | **0.00364** | **2e-05** | **0** | **0** | **0.00515** | **2e-05** | **0** | **0** |
| Median | **0.00728** | **0.00011** | **0** | **0** | **0.01008** | **0.00014** | **0** | **0** |
| Mean | **0.00808** | **0.00049** | **0** | **0** | **0.01014** | **0.00065** | **0** | **0** |
| 3rd Quart | **0.01162** | **0.00046** | **0** | **0** | **0.01477** | **0.00061** | **0** | **0** |
| Max | **0.02881** | **0.01397** | **0** | **0** | **0.02401** | **0.01796** | **0** | **0** |

**Posthoc tests using genus-level full dataset**

Table S13: Summary statistics for pairwise corrected p-value distributions for phyANOVA using all 10000 genus-level dataset. Values were rounded to the third decimal place.

| Epsilon | Threshold | State 1 | State 2 | Min | 1st Quart | Median | Mean | 3rd Quart | Max |
|---|---|---|---|---|---|---|---|---|---|
| 0.0 | 50% | AM | EM | 0.873 | 1.000 | 1.000 | 1.000 | 1.000 | 1.000 |
| 0.0 | 50% | AM | MIX | 0.006 | 0.384 | 1.000 | 0.719 | 1.000 | 1.000 |
| 0.0 | 50% | EM | MIX | 0.008 | 1.000 | 1.000 | 0.957 | 1.000 | 1.000 |
| 0.0 | 50% | AM | NM | 0.618 | 1.000 | 1.000 | 1.000 | 1.000 | 1.000 |
| 0.0 | 50% | EM | NM | 0.886 | 1.000 | 1.000 | 1.000 | 1.000 | 1.000 |
| 0.0 | 50% | MIX | NM | 0.006 | 0.509 | 1.000 | 0.765 | 1.000 | 1.000 |
| 0.0 | 60% | AM | EM | 0.820 | 1.000 | 1.000 | 1.000 | 1.000 | 1.000 |
| 0.0 | 60% | AM | MIX | 0.006 | 0.006 | 0.024 | 0.080 | 0.078 | 1.000 |
| 0.0 | 60% | EM | MIX | 0.180 | 1.000 | 1.000 | 0.989 | 1.000 | 1.000 |
| 0.0 | 60% | AM | NM | 0.829 | 1.000 | 1.000 | 1.000 | 1.000 | 1.000 |
| 0.0 | 60% | EM | NM | 0.720 | 1.000 | 1.000 | 1.000 | 1.000 | 1.000 |
| 0.0 | 60% | MIX | NM | 0.006 | 0.010 | 0.030 | 0.116 | 0.110 | 1.000 |
| 0.0 | 80% | AM | EM | 0.726 | 1.000 | 1.000 | 0.997 | 1.000 | 1.000 |
| 0.0 | 80% | AM | MIX | 0.006 | 0.006 | 0.006 | 0.006 | 0.006 | 0.018 |
| 0.0 | 80% | EM | MIX | 0.740 | 1.000 | 1.000 | 0.997 | 1.000 | 1.000 |
| 0.0 | 80% | AM | NM | 0.712 | 1.000 | 1.000 | 0.997 | 1.000 | 1.000 |
| 0.0 | 80% | EM | NM | 0.556 | 1.000 | 1.000 | 0.988 | 1.000 | 1.000 |
| 0.0 | 80% | MIX | NM | 0.006 | 0.006 | 0.006 | 0.006 | 0.006 | 0.025 |
| 0.0 | 100% | AM | EM | 0.228 | 0.396 | 0.436 | 0.431 | 0.472 | 0.608 |
| 0.0 | 100% | AM | MIX | 0.006 | 0.006 | 0.006 | 0.006 | 0.006 | 0.006 |
| 0.0 | 100% | EM | MIX | 0.254 | 0.560 | 0.606 | 0.578 | 0.634 | 0.750 |
| 0.0 | 100% | AM | NM | 0.511 | 0.616 | 0.645 | 0.648 | 0.680 | 0.853 |
| 0.0 | 100% | EM | NM | 0.220 | 0.396 | 0.436 | 0.431 | 0.472 | 0.608 |
| 0.0 | 100% | MIX | NM | 0.006 | 0.006 | 0.006 | 0.006 | 0.006 | 0.006 |
| 0.9 | 50% | AM | EM | 1.000 | 1.000 | 1.000 | 1.000 | 1.000 | 1.000 |
| 0.9 | 50% | AM | MIX | 0.006 | 0.534 | 1.000 | 0.778 | 1.000 | 1.000 |
| 0.9 | 50% | EM | MIX | 0.008 | 1.000 | 1.000 | 0.939 | 1.000 | 1.000 |
| 0.9 | 50% | AM | NM | 1.000 | 1.000 | 1.000 | 1.000 | 1.000 | 1.000 |
| 0.9 | 50% | EM | NM | 1.000 | 1.000 | 1.000 | 1.000 | 1.000 | 1.000 |
| 0.9 | 50% | MIX | NM | 0.006 | 0.540 | 1.000 | 0.777 | 1.000 | 1.000 |
| 0.9 | 60% | AM | EM | 0.696 | 1.000 | 1.000 | 1.000 | 1.000 | 1.000 |
| 0.9 | 60% | AM | MIX | 0.006 | 0.012 | 0.040 | 0.118 | 0.130 | 1.000 |
| 0.9 | 60% | EM | MIX | 0.100 | 1.000 | 1.000 | 0.956 | 1.000 | 1.000 |
| 0.9 | 60% | AM | NM | 0.624 | 1.000 | 1.000 | 1.000 | 1.000 | 1.000 |
| 0.9 | 60% | EM | NM | 0.696 | 1.000 | 1.000 | 1.000 | 1.000 | 1.000 |
| 0.9 | 60% | MIX | NM | 0.006 | 0.006 | 0.024 | 0.098 | 0.085 | 1.000 |
| 0.9 | 80% | AM | EM | 0.654 | 1.000 | 1.000 | 0.982 | 1.000 | 1.000 |
| 0.9 | 80% | AM | MIX | 0.006 | 0.006 | 0.006 | 0.007 | 0.006 | 0.045 |
| 0.9 | 80% | EM | MIX | 0.484 | 1.000 | 1.000 | 0.980 | 1.000 | 1.000 |
| 0.9 | 80% | AM | NM | 0.504 | 0.996 | 1.000 | 0.968 | 1.000 | 1.000 |
| 0.9 | 80% | EM | NM | 0.560 | 1.000 | 1.000 | 0.973 | 1.000 | 1.000 |
| 0.9 | 80% | MIX | NM | 0.006 | 0.006 | 0.006 | 0.006 | 0.006 | 0.035 |
| 0.9 | 100% | AM | EM | 0.315 | 0.460 | 0.492 | 0.493 | 0.524 | 0.680 |
| 0.9 | 100% | AM | MIX | 0.006 | 0.006 | 0.006 | 0.006 | 0.006 | 0.006 |
| 0.9 | 100% | EM | MIX | 0.192 | 0.486 | 0.512 | 0.509 | 0.538 | 0.680 |
| 0.9 | 100% | AM | NM | 0.486 | 0.562 | 0.585 | 0.586 | 0.608 | 0.721 |
| 0.9 | 100% | EM | NM | 0.312 | 0.460 | 0.492 | 0.492 | 0.524 | 0.680 |
| 0.9 | 100% | MIX | NM | 0.006 | 0.006 | 0.006 | 0.006 | 0.006 | 0.006 |

Table S14: Summary statistics for pairwise corrected p-value distributions for standard ANOVA using all 10000 genus-level dataset. Values were rounded to the third decimal place.

| Epsilon | Threshold | State 1 | State 2 | Min | 1st Quart | Median | Mean | 3rd Quart | Max |
|---|---|---|---|---|---|---|---|---|---|
| 0.0 | 50% | AM | EM | 0.399 | 0.609 | 0.706 | 0.694 | 0.775 | 0.887 |
| 0.0 | 50% | AM | MIX | 0.000 | 0.235 | 0.537 | 0.530 | 0.828 | 1.000 |
| 0.0 | 50% | EM | MIX | 0.005 | 0.825 | 0.959 | 0.868 | 0.995 | 1.000 |
| 0.0 | 50% | AM | NM | 0.079 | 0.664 | 0.826 | 0.784 | 0.939 | 1.000 |
| 0.0 | 50% | EM | NM | 0.405 | 0.847 | 0.911 | 0.892 | 0.954 | 1.000 |
| 0.0 | 50% | MIX | NM | 0.000 | 0.387 | 0.740 | 0.653 | 0.953 | 1.000 |
| 0.0 | 60% | AM | EM | 0.323 | 0.610 | 0.690 | 0.676 | 0.761 | 0.865 |
| 0.0 | 60% | AM | MIX | 0.000 | 0.001 | 0.003 | 0.021 | 0.017 | 0.781 |
| 0.0 | 60% | EM | MIX | 0.057 | 0.614 | 0.776 | 0.740 | 0.900 | 1.000 |
| 0.0 | 60% | AM | NM | 0.297 | 0.951 | 0.990 | 0.958 | 0.999 | 1.000 |
| 0.0 | 60% | EM | NM | 0.147 | 0.674 | 0.772 | 0.755 | 0.849 | 0.996 |
| 0.0 | 60% | MIX | NM | 0.000 | 0.005 | 0.023 | 0.080 | 0.087 | 0.998 |
| 0.0 | 80% | AM | EM | 0.259 | 0.540 | 0.559 | 0.575 | 0.626 | 0.713 |
| 0.0 | 80% | AM | MIX | 0.000 | 0.000 | 0.000 | 0.000 | 0.000 | 0.000 |
| 0.0 | 80% | EM | MIX | 0.241 | 0.627 | 0.723 | 0.705 | 0.796 | 0.954 |
| 0.0 | 80% | AM | NM | 0.340 | 0.695 | 0.800 | 0.787 | 0.892 | 1.000 |
| 0.0 | 80% | EM | NM | 0.142 | 0.327 | 0.379 | 0.388 | 0.441 | 0.732 |
| 0.0 | 80% | MIX | NM | 0.000 | 0.000 | 0.000 | 0.000 | 0.000 | 0.007 |
| 0.0 | 100% | AM | EM | 0.073 | 0.095 | 0.097 | 0.096 | 0.099 | 0.114 |
| 0.0 | 100% | AM | MIX | 0.000 | 0.000 | 0.000 | 0.000 | 0.000 | 0.000 |
| 0.0 | 100% | EM | MIX | 0.133 | 0.364 | 0.387 | 0.365 | 0.404 | 0.427 |
| 0.0 | 100% | AM | NM | 0.860 | 0.911 | 0.933 | 0.929 | 0.951 | 0.995 |
| 0.0 | 100% | EM | NM | 0.048 | 0.070 | 0.080 | 0.079 | 0.088 | 0.128 |
| 0.0 | 100% | MIX | NM | 0.000 | 0.000 | 0.000 | 0.000 | 0.000 | 0.000 |
| 0.9 | 50% | AM | EM | 0.669 | 0.887 | 0.922 | 0.914 | 0.965 | 0.998 |
| 0.9 | 50% | AM | MIX | 0.000 | 0.315 | 0.649 | 0.602 | 0.910 | 1.000 |
| 0.9 | 50% | EM | MIX | 0.003 | 0.755 | 0.939 | 0.834 | 0.993 | 1.000 |
| 0.9 | 50% | AM | NM | 0.241 | 0.926 | 0.982 | 0.943 | 0.998 | 1.000 |
| 0.9 | 50% | EM | NM | 0.428 | 0.930 | 0.971 | 0.951 | 0.991 | 1.000 |
| 0.9 | 50% | MIX | NM | 0.000 | 0.385 | 0.740 | 0.652 | 0.954 | 1.000 |
| 0.9 | 60% | AM | EM | 0.604 | 0.863 | 0.926 | 0.901 | 0.951 | 0.995 |
| 0.9 | 60% | AM | MIX | 0.000 | 0.001 | 0.008 | 0.039 | 0.036 | 0.954 |
| 0.9 | 60% | EM | MIX | 0.020 | 0.436 | 0.623 | 0.608 | 0.795 | 1.000 |
| 0.9 | 60% | AM | NM | 0.180 | 0.882 | 0.968 | 0.915 | 0.995 | 1.000 |
| 0.9 | 60% | EM | NM | 0.191 | 0.765 | 0.854 | 0.834 | 0.923 | 1.000 |
| 0.9 | 60% | MIX | NM | 0.000 | 0.002 | 0.014 | 0.063 | 0.062 | 0.985 |
| 0.9 | 80% | AM | EM | 0.535 | 0.766 | 0.785 | 0.822 | 0.878 | 0.966 |
| 0.9 | 80% | AM | MIX | 0.000 | 0.000 | 0.000 | 0.000 | 0.000 | 0.000 |
| 0.9 | 80% | EM | MIX | 0.130 | 0.508 | 0.638 | 0.618 | 0.745 | 0.925 |
| 0.9 | 80% | AM | NM | 0.118 | 0.346 | 0.439 | 0.457 | 0.554 | 0.963 |
| 0.9 | 80% | EM | NM | 0.196 | 0.382 | 0.445 | 0.454 | 0.517 | 0.820 |
| 0.9 | 80% | MIX | NM | 0.000 | 0.000 | 0.000 | 0.000 | 0.000 | 0.005 |
| 0.9 | 100% | AM | EM | 0.113 | 0.125 | 0.128 | 0.136 | 0.131 | 0.224 |
| 0.9 | 100% | AM | MIX | 0.000 | 0.000 | 0.000 | 0.000 | 0.000 | 0.000 |
| 0.9 | 100% | EM | MIX | 0.036 | 0.270 | 0.289 | 0.258 | 0.304 | 0.322 |
| 0.9 | 100% | AM | NM | 0.830 | 0.875 | 0.902 | 0.897 | 0.906 | 0.968 |
| 0.9 | 100% | EM | NM | 0.066 | 0.087 | 0.094 | 0.099 | 0.106 | 0.188 |
| 0.9 | 100% | MIX | NM | 0.000 | 0.000 | 0.000 | 0.000 | 0.000 | 0.000 |

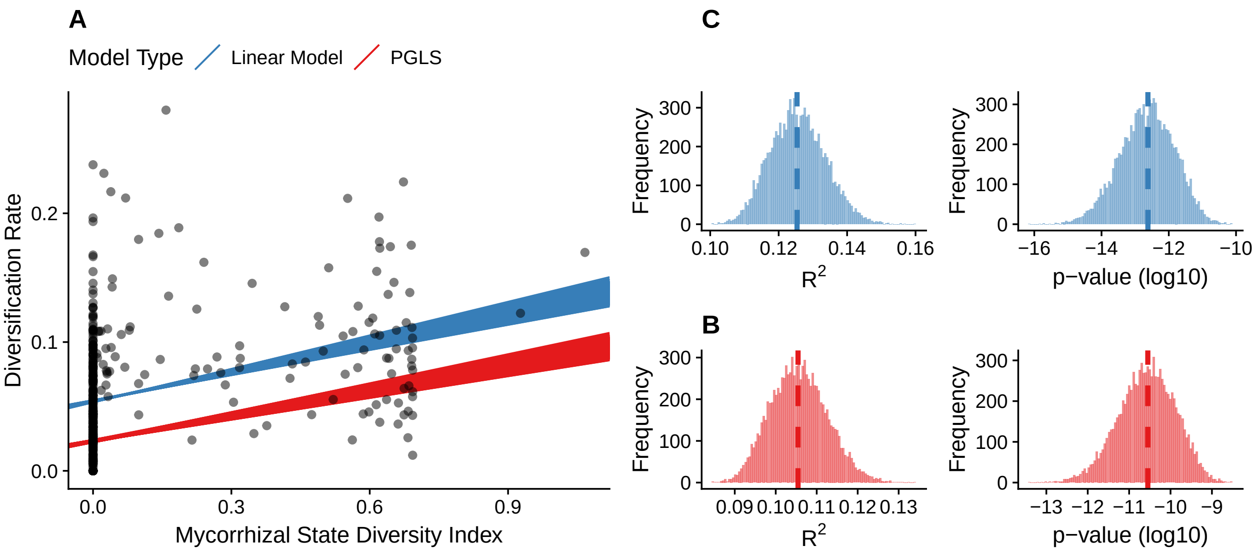

Figure S17: Scatterplots showing a randomly selected dataset replicate (out of 10000) to show the relationship between mycorrhizal state diversity index and diversification rates estimated with $\epsilon$ (relative extinction fraction) $= 0.9$. The red and blue lines indicate the linear models and phylogenetic generalized least squares (PGLS) fit, respectively, for 1000 datasets of seed plant mycorrhizal states. Panels b and c show the frequency distribution of R2 and p-values of (b) linear models (blue bars) and (c) PGLS (red bars), dashed lines show the median for each distribution.

Table S15: Summary statistics for p-value and $R^2$ for the phylogenetic and parametric regressions using all 10000 genus-level datasets.

| | PGLS | | | | LM | | | |
|---|---|---|---|---|---|---|---|---|
| | e = 0 | | e = 0.9 | | e = 0 | | e = 0.9 | |
| Quantile | p-value | $R^2$ | p-value | $R^2$ | p-value | $R^2$ | p-value | $R^2$ |
| Min | 0 | 0.085 | 0 | 0.081 | 0 | 0.101 | 0 | 0.092 |
| 1st Quart | 0 | 0.101 | 0 | 0.100 | 0 | 0.120 | 0 | 0.113 |
| Median | 0 | 0.105 | 0 | 0.105 | 0 | 0.125 | 0 | 0.119 |
| Mean | 0 | 0.106 | 0 | 0.105 | 0 | 0.126 | 0 | 0.119 |
| 3rd Quart | 0 | 0.110 | 0 | 0.110 | 0 | 0.131 | 0 | 0.124 |
| Max | 0 | 0.134 | 0 | 0.135 | 0 | 0.160 | 0 | 0.153 |

### Appendix S6 - Replicating main analyses with tree from Harris and Davies (2016)

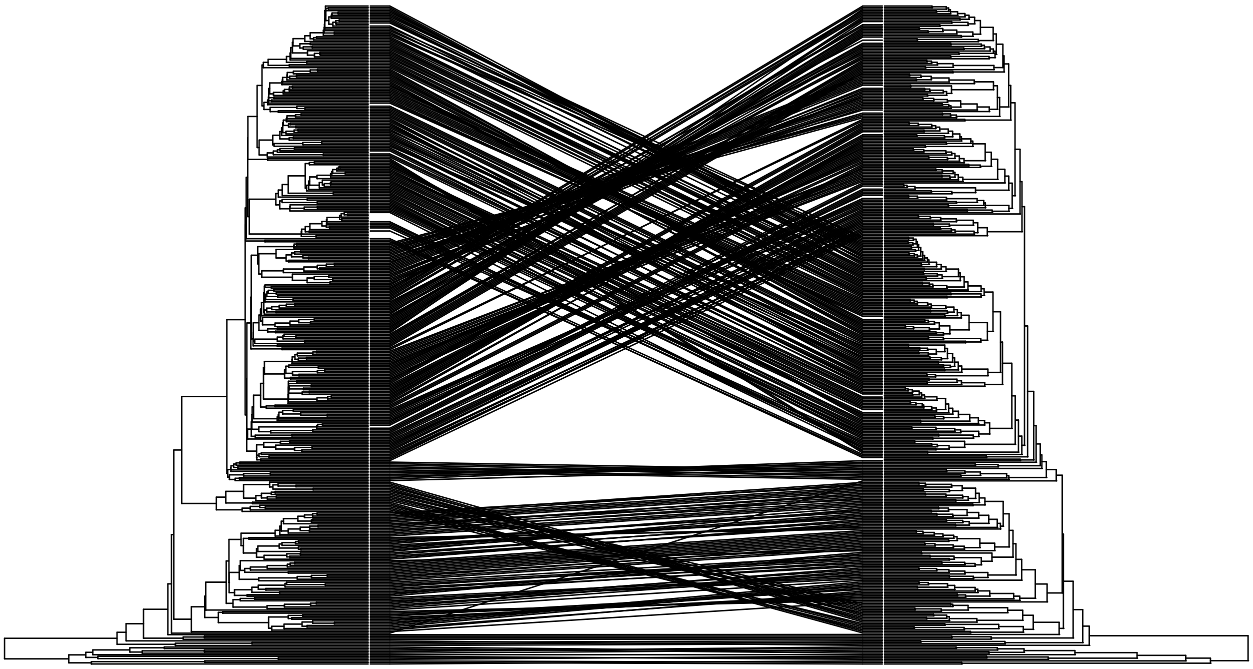

Figure S18: Association between placement of families in both Zanne's (left) and Harris and Davies' (right) phylogenies, indicating differences in the topologies.

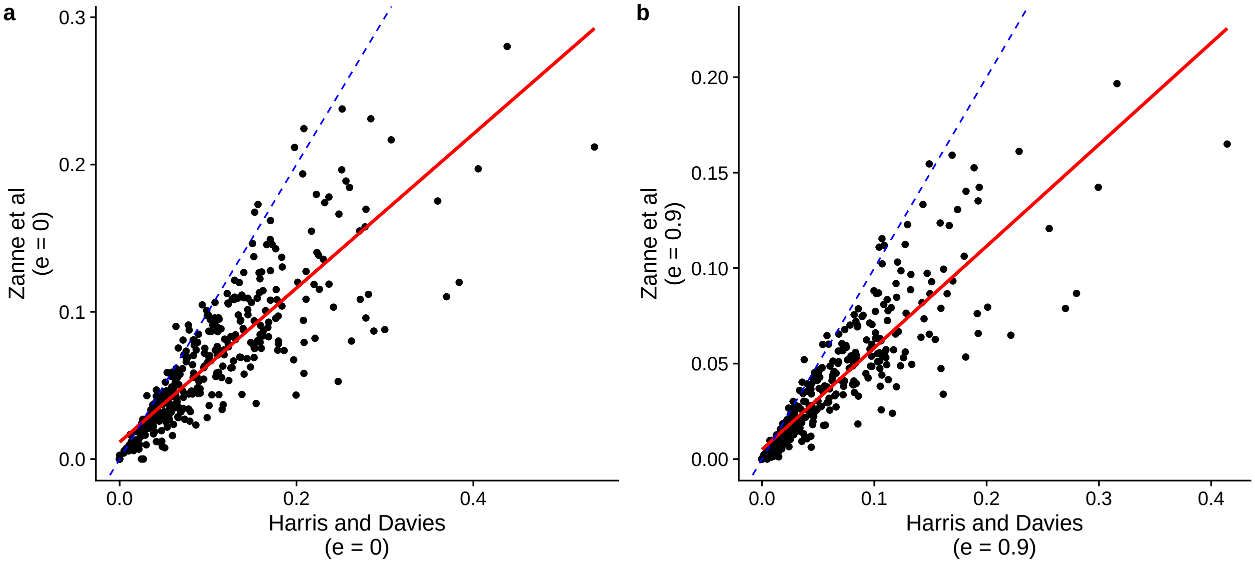

Figure S19: Relationship between net diversification calculated in our study and with the rates from Harris and Davies (2016) The rates were estimated by the authors using the same source for species richness as the main analysis, and the stem ages were obtained from their phylogeny. a: $\epsilon = 0$; b: $\epsilon = 0.9$. Red lines: LM fit; blue dashed lines: identity fit.

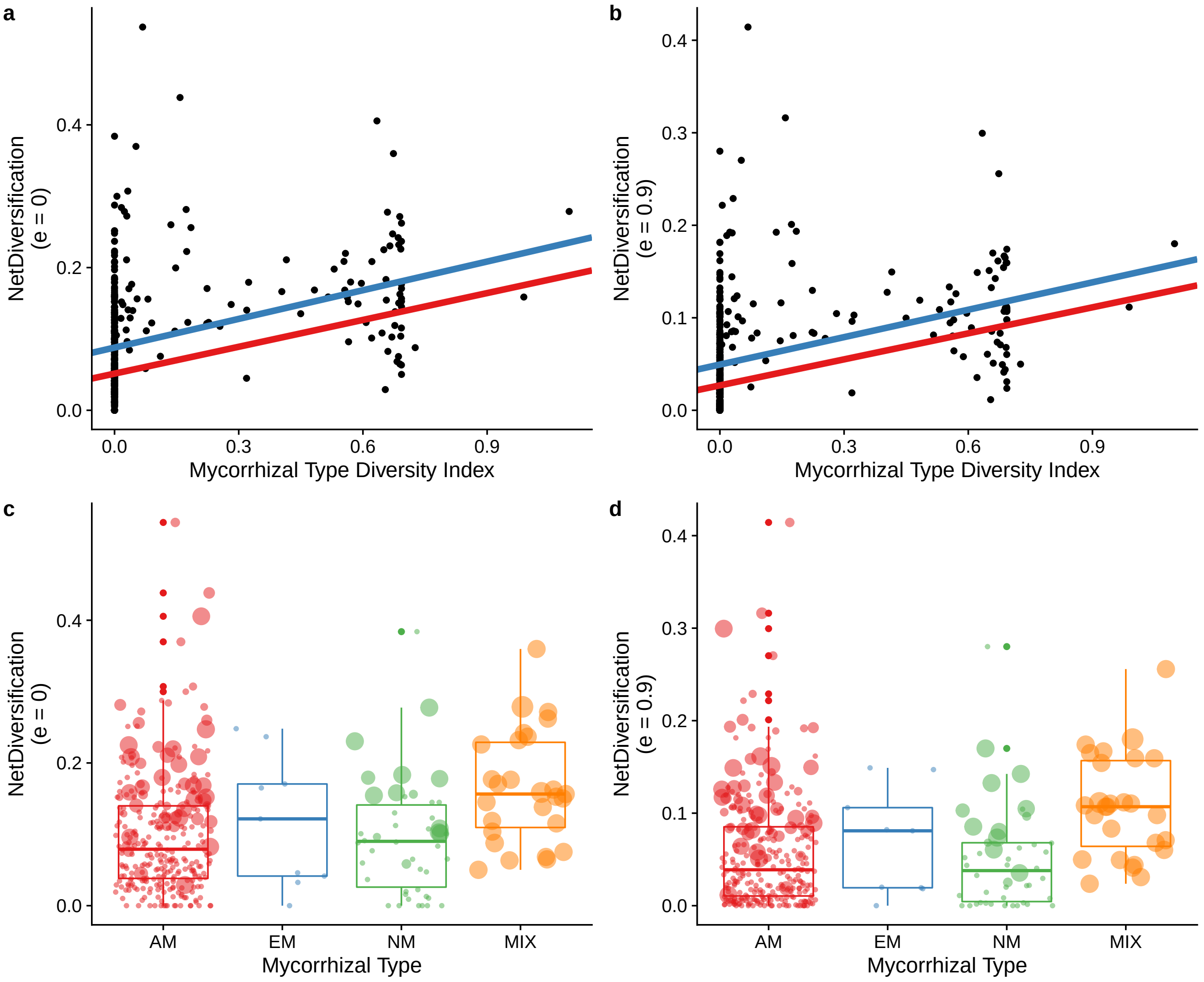

Figure S20: Main analyses replicated using Harris and Davies (2016) phylogeny. The rates were estimated by the authors using the same source for species richness as the main analysis, and the stem ages were obtained from their phylogeny. The models were tested using only the empirical dataset, with no randomization. Panels a and c: $\epsilon = 0$; panels b and d: $\epsilon = 0.9$. Red lines: PGLS fit; blue dashed lines: LM fit.

Table S16: Summary statistics for linear models and ANOVA using Harris and Davies (2016) phylogeny.

| | e = 0 | | | e = 0.9 | | |
|---|---|---|---|---|---|---|
| | Regression | | ANOVA | Regression | | ANOVA |
| Model | p-value | $R^2$ | p-value | p-value | $R^2$ | p-value |
| Phylogenetic | **2.724e-13** | 0.135 | **0.0005** | **3.581e-13** | 0.134 | **0.0003** |
| Linear | **1.345e-14** | 0.146 | **0.019** | **1.228e-14** | 0.147 | **0.02** |

#### Appendix S7 - Phylogenetic signal of diversification rates, age, and richness

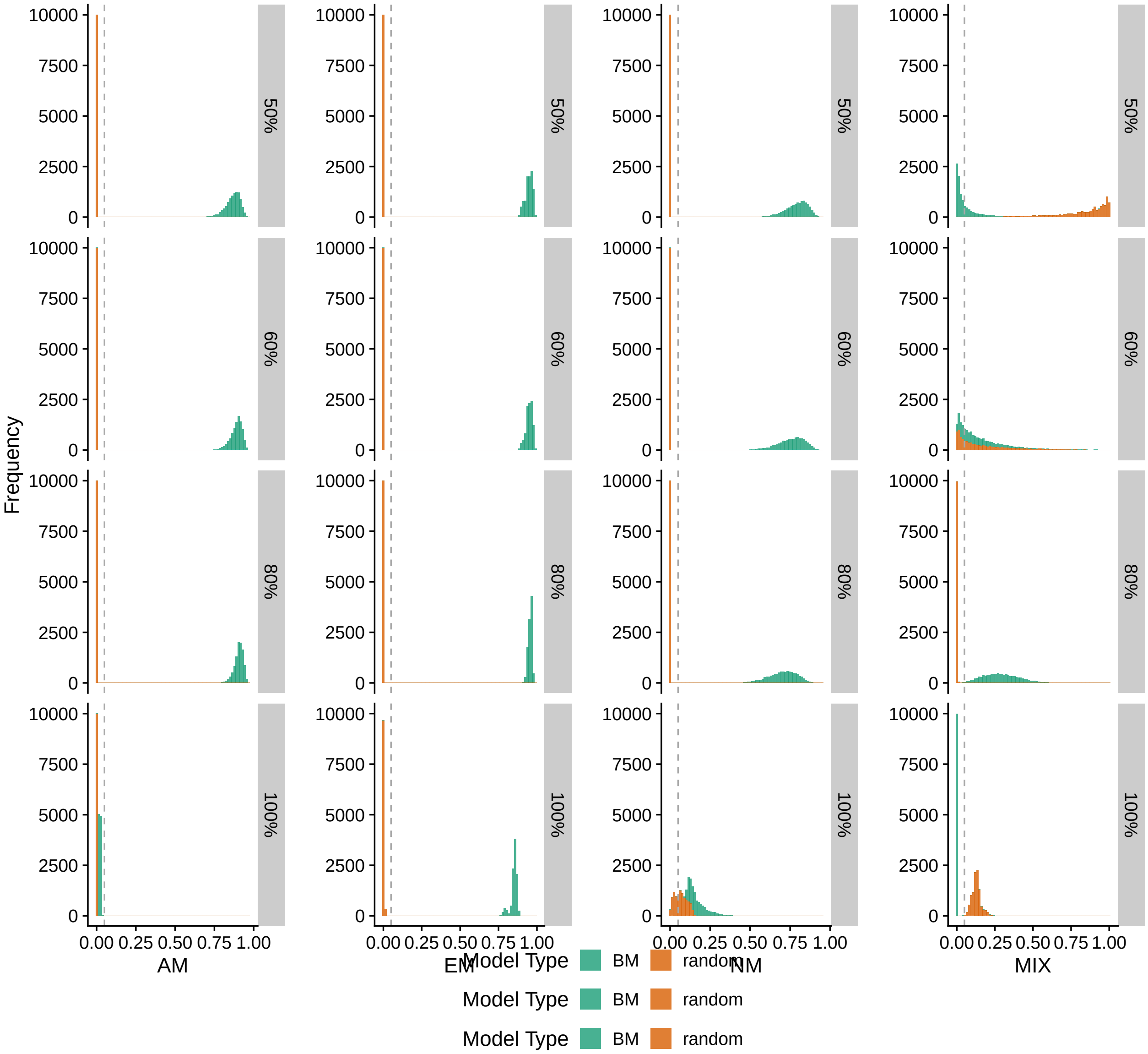

Figure S21: P-value distribution of phylogenetic signal of each mycorrhizal state using different thresholds for classification. Colour indicates each model to which the phylogenetic structure of mycorrhizal type is compared, and vertical dashed line indicates the value of 0.05

Table S17: Phylogenetic signal of the four response variables using the genus-level dataset including all families. Significant values are highlighted in bold.

| Variable | Lambda |
|---|---|
| r (epsilon = 0) | **0.474** |
| r (epsilon = 0.9) | **0.357** |
| Stem Age | **1** |
| Richness | 1e-06 |

### Appendix S8 - Percentage of MIX species

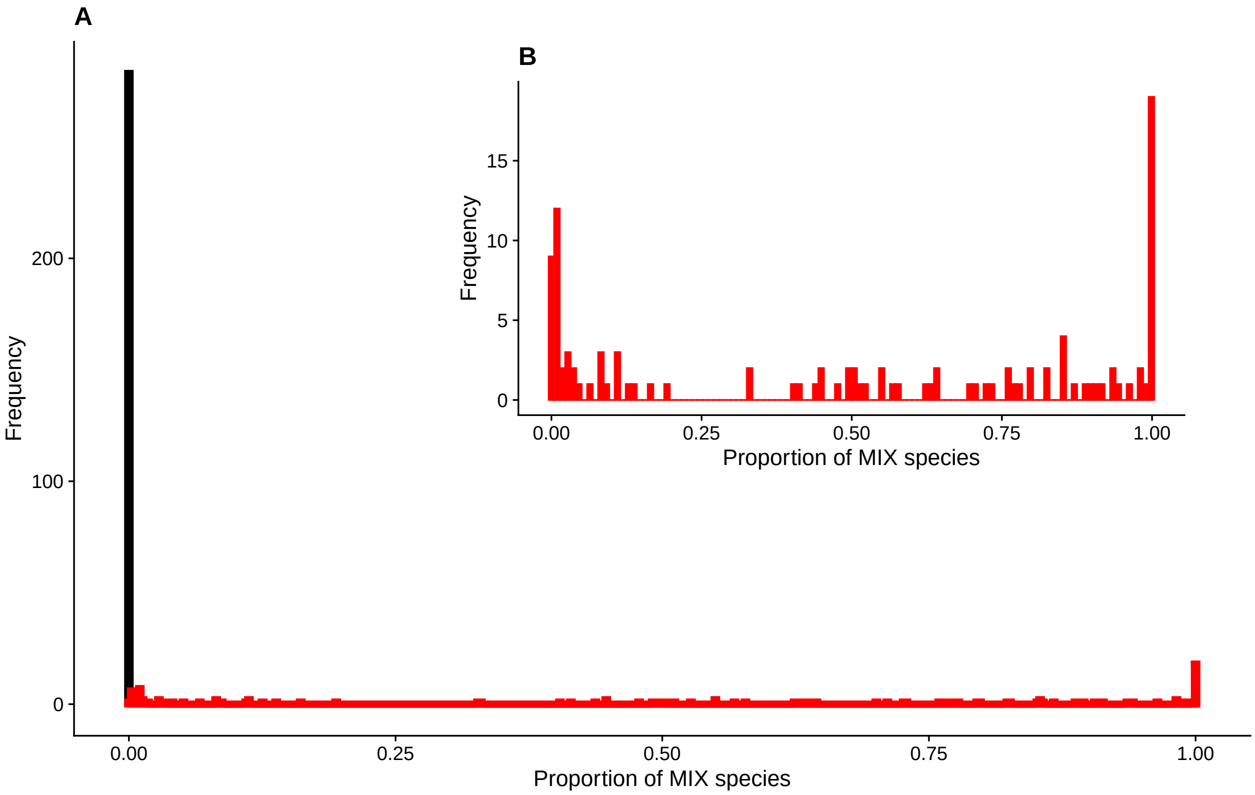

Figure S22: Distribution of percentage of species in each family that are classified as MIX. Zoom panel indicates the same distribution but excluding families without any MIX species.
